# A highly contiguous hexaploid wheat genome assembly facilitates analysis of 1RS translocation and mining of a new adult plant resistance locus to yellow rust disease

**DOI:** 10.1101/2023.12.30.573687

**Authors:** Guangwei Li, Yan Ren, Yuxin Yang, Shulin Chen, Jizhou Zheng, Xiaoqing Zhang, Mengen Chen, Xiaonan Sun, Chunlei Lv, Xiaode Li, Yujia Li, Chunhao Dong, Jianwei Tang, Zhenpu Huang, Yanyan Peng, Dengbin Gu, Zhiyong Wang, Hongyuan Zheng, Cuilan Shi, Guozhang Kang, Tiancun Zheng, Feng Chen, Daowen Wang, Kunpu Zhang, Guihong Yin

## Abstract

High-quality genome information is essential for efficiently deciphering and improving crop traits. Here we report a highly contiguous hexaploid genome assembly for the key wheat breeding parent Zhou8425B, an elite 1BL/1RS translocation line with durable adult plant resistance (APR) against rust diseases. By using HiFi and Hi-C sequencing reads, a 14.75 Gb genome assembly, with contig N50 and scaffold N50 values reaching 70.94 and 735.11 Mb, respectively, was developed. Comparison with 16 previously sequenced common wheat cultivars revealed unique chromosomal structural features in Zhou8425B. Notably, the 1RS translocation in Zhou8425B was apparently longer and carried more genes encoding AP2/ERF-ERF and B3 transcription factors relative to its counterpart in several genome sequenced 1BL/1RS varieties and rye lines. Aided by Zhou8425B genome assembly, a new APR locus (i.e., *YrZH3B*) against yellow rust (YR) disease was finely mapped to a 1 - 2 Mb interval on chromosome 3BS. Analysis with 212 Zhou8425B derivative varieties showed that pyramiding of *YrZH3B* with two other APR loci (*YrZH22* and *YrZH84*) significantly decreased YR severity and enhanced grain yield, with triple combination (*YrZH3B/YrZH22/YrZH84*) having the highest effects. Our data demonstrate the high value of Zhou8425B assembly in studying wheat genome and agronomically important genes.

## Introduction

Common wheat (*Triticum aestivum*, AABBDD, 2n = 6x = 42) is staple food for over 35% of the world population. Its efficient improvement is essential for maintaining the food and nutrition security of an ever-increasing global population in the era of dwindling agricultural resources and intensifying climate change. However, it is a challenging task to improve wheat traits efficiently as they are controlled by polygenes and affected by environmental factors. Fortunately, the advancement of plant genomics and genome editing has largely quickened the pace of dissecting and enhancing crop traits (*1, 2*). Since 2014, genome sequence has been reported for an increasing number of wheat accessions and its relatives (*3–5*). More recently, there is a growing interest internationally to sequence elite wheat germplasms to accelerate the mining of genes controlling important agronomic traits (*6–9*).

Zhou8425B is a semidwarf 1BL/1RS translocation line and exhibits superior traits including durable adult plant resistance (APR) to yellow rust (YR) and leaf rust (LR) diseases (*10*). Using Zhou8425B and derivative varieties, four major APR loci against YR (*YrZH22* and *YrZH84*) or LR (*LrZH22* and *LrZH84*) have been mapped on 1B (*LrZH84*), 2B (*LrZH22*), 4B (*YrZH22*), and 7B (*YrZH84*) chromosomes (*11–15*). *LrZH22* (*Lr13*) was recently isolated by map-based cloning, and found to encode a nucleotide-binding leucine-rich repeat (NLR) protein with coiled-coil, nucleotide-binding site, and leucine-rich repeat domains (*16*). However, the molecular identities of *LrZH84*, *YrZH22* and *YrZH84* remain obscure at present. In addition to possessing APR to rust diseases, Zhou8425B also shows elite yield related traits (such as higher grain number per spike and larger grains) and high tolerance to abiotic stresses (e.g., drought) (*17–22*). More than 70 important quantitative trait loci (QTLs) linked to the superior stress tolerance and yield related traits of Zhou8425B, with the percentage of variance explained (PVE) ≥ 10%, have been detected over the last 20 years (*11–15*, *17–22*), but none of them have been molecularly cloned to date. Lack of genome sequence information may have contributed to the slow progress in characterizing the valuable genes carried by Zhou8425B.

Owing to the outstanding agronomic traits, Zhou8425B has been used as a founder parent in wheat breeding in China since its release in 1988 (*23–26*). To aid characterization of the valuable genes in Zhou8425B and genomics assisted breeding in wheat, we developed a highly contiguous genome assembly for Zhou8425B using PacBio HiFi sequencing platform and high throughput chromosome conformation capture sequencing (Hi-C) technology. The quality of Zhou8425B genome assembly, as judged by contig and scaffold N50 values and several other parameters, was better than that of the hexaploid wheat cultivars sequenced before (*3–9*). The unique chromosomal translocations and highly tandemly duplicated gene clusters in Zhou8425B were identified by comparison with previously sequenced hexaploid wheat cultivars. Interestingly, the 1RS translocation chromosome arm in Zhou8425B was longer and harbored more genes predicted to encode AP2/ERF-ERF or B3 transcription factors (TFs) relative to that in the two hexaploid wheat varieties (Kenong 9204 and Aikang 58) and two diploid rye lines (Lo7 and Weining) with whole genome sequence information (*8, 9, 27, 28*). Finally, Zhou8425B genome assembly was employed to accelerate the mapping of a new APR locus (tentatively named as *YrZH3BS*) to a 1 - 2 Mb interval on chromosome 3BS. The genetic effects of *YrZH3BS* as well as its combinations with *YrZH22* and *YrZH84* on decreasing YR disease severity and enhancing grain yield were validated by analyzing 212 Zhou8425B derivative varieties. The development and analysis of Zhou8425B genome assembly has increased the understanding of wheat genome diversities and agronomic traits and will contribute importantly to genomics-assisted wheat breeding in the future.

## Results

### Pedigree, agronomic traits, and breeding application of Zhou8425B

Zhou8425B was developed in China from 1978 to 1988 in order to make use of the elite genes carried by 1RS translocation (*23, 24*). Its pedigree involved the hexaploid triticale line Guangmai 74 (AABBRR) and a number of common wheat varieties originated in European countries (Former Soviet Union, Italy, and Britain) and China, with the breeding process encompassing multiple rounds of crossing and backcrossing and one times of γ irradiation treatment of dry seeds (Fig. 1A). It was semidwarf (∼70 cM) and had relatively longer spike (∼13.1 cM), higher number of spikelets (∼13.1), and heavier thousand grain weight (∼ 50 g) (Fig. 1B-D). Previous studies have identified four major APR loci against YR (*YrZH22* and *YrZH84*) and LR (*LrZH22*/*Lr13* and *LrZH84*) using Zhou8425B or its early-generation derivative cultivar Zhoumai 22 as experimental materials (*11–15*). Additionally, Zhou8425B carried a large number of elite QTL alleles regulating grain yield traits, such as those associated with spike length or grain number per spike (e.g., *QSl.nafu-6A.2*, *QSl.nafu-7A*, and *QGns.nafu-2B*) (*17, 20, 29*), as well as four dwarf genes (*Rht1*, *Rht2*, *Rht8*, and *Rht24*) (*30*).

**Figure 1.**
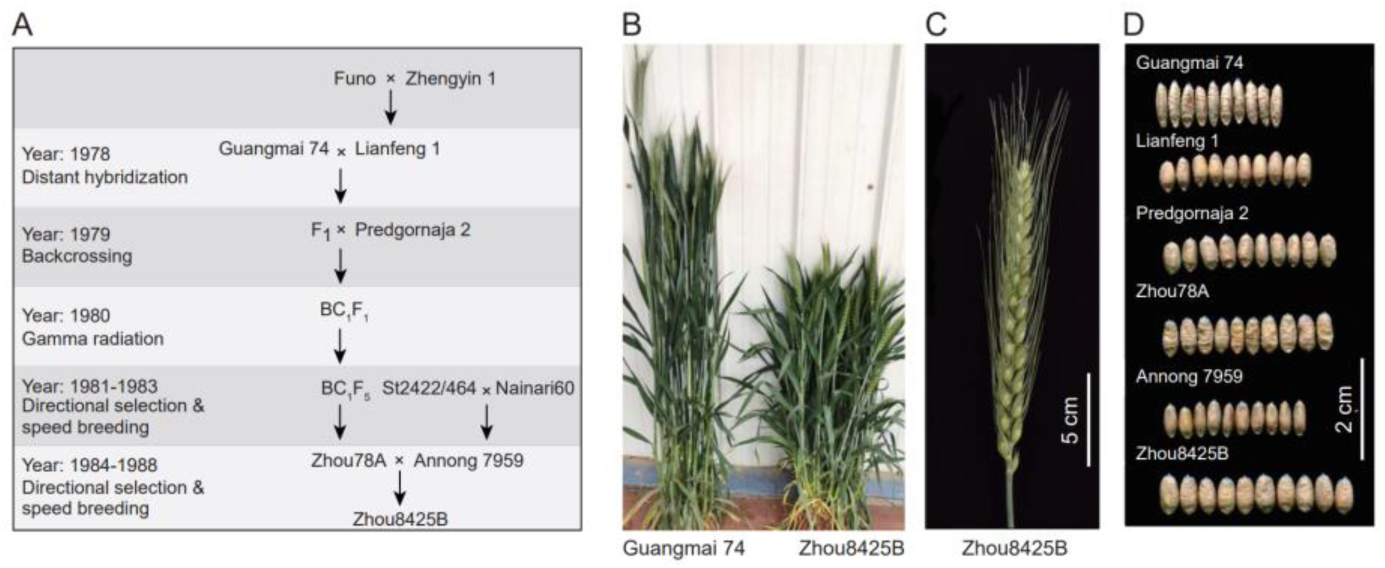
Breeding of Zhou8425B and its agronomic traits. **A.** The timeline, wheat materials, and genetic crosses involved in developing Zhou8425B. Guangmai 74 is a hexaploid triticale, whereas the other lines are hexaploid common wheat. A γ-ray irradiation treatment was applied to the BC_1_F_1_ seeds in 1980. **B.** Comparison of plant height between Guangmai 74 and Zhou8425B. **C.** A typical spike of Zhou8425B plants grown under field conditions. **D.** Comparison of grain morphology and size between Zhou8425B and the five different wheat materials used during the breeding of Zhou8425B.

To better understand the origin of important trait genes in Zhou8425B, we revealed differential presence of four dwarf genes, the photoperiod insensitive gene *Ppd-D1a*, five APR loci including the newly mined *YrZH3BS* locus against YR disease (see below), and seven grain weight associated genes (*TaCwi-A1*, *TaSus1-7A*, *TaSus1-7B*, *TaSus2-2A*, *TaGS-D1*, *TaGS5-A1*, and *TaGW2-6B*) in the genetic materials used in the breeding of Zhou8425B by molecular marker analysis (Table 1). Most of the examined genes were effectively pyramided in Zhou78A, the immediate maternal parent of Zhou8425B, with the paternal parent Annong 7959 providing *Rht24* and the elite alleles of *TaCwi-A1* and *TaGS5-A1* (Table 1; Fig. 1A).

**Table 1.** Detection of 17 agronomic trait genes in Zhou8425B and the wheat lines used during Zhou8425B development by DNA marker analysis.

| Gene | Guangmai 74 | Lianfeng 1 | Predgornaja 2 | Zhou78A | Annong 7959 | Zhou8425B |
| --- | --- | --- | --- | --- | --- | --- |
| *Rht1* | + | + | + | + | + | + |
| *Rht2* | + | + | + | + | + | + |
| *Rht8* | - | + | + | + | - | + |
| *Rht24* | - | + | - | - | + | + |
| *Ppd-D1a* | - | + | + | + | + | + |
| *YrZH84* | + | - | - | + | - | + |
| *YrZH22* | + | - | + | + | - | + |
| *YrZH3BS* | - | + | - | + | + | + |
| *LrZH84* | + | - | + | + | - | + |
| *Lr13* | + | - | - | + | - | + |
| *TaCwi-A1* | + | + | - | - | + | + |
| *TaSus1-7A* | + | + | + | + | + | + |
| *TaSus1-7B* | - | + | + | + | + | + |
| *TaSus2-2A* | + | + | + | + | + | + |
| *TaGS-D1* | + | + | + | + | + | + |
| *TaGS5-A1* | + | + | - | - | + | + |
| *TaGW2-6B* | - | + | - | + | - | + |

+, presence; -, absence.

Since 1988, Zhou8425B and its early-generation descendants (Zhoumai 11, 12, 13, 16, 17 & 22 and Aikang 58) have been extensively used as wheat breeding parents in China, with a total of 839 derivative cultivars passed provincial and/or national certification by 2023 (Table S1). Many derivative varieties, such Zhoumai 16, Zhoumai 22, Aikang 58, Zhengmai 7698, and Bainong 207, have gained wide applications in wheat production in the major winter wheat production zone (i.e., the Yellow-Huai River Valley region) of China (*31, 32*).

### Characteristics of Zhou8425B genome assembly

As the first step of constructing Zhou8425B genome assembly, we checked its chromosomal number and 1RS translocation using fluorescent *in situ* hybridization assay. A total of 42 somatic chromosomes were consistently found in root tip cells, with one pair of 1BL/1RS translocation chromosomes clearly identified based on fluorescent signals produced using the hybridization probes pSC119.2 and pTa535 (Fig. 2A). Subsequently, a 14.75 Gb genome assembly, with contig and scaffold N50 reaching 70.94 and 735.11 Mb, respectively (Table 2), was constructed by combing HiFi (48X genome coverage) and Hi-C (124X genome depth) sequencing reads (Tables S2 and S3). Approximately 98.44% (14.52 Gb) of the assembly were anchored onto 21 chromosomes based on the chromatin interactions revealed by Hi-C analysis (Fig. S1); the remaining 0.23 Gb unanchored sequences were mainly rDNA or other repeats.

**Figure 2.**
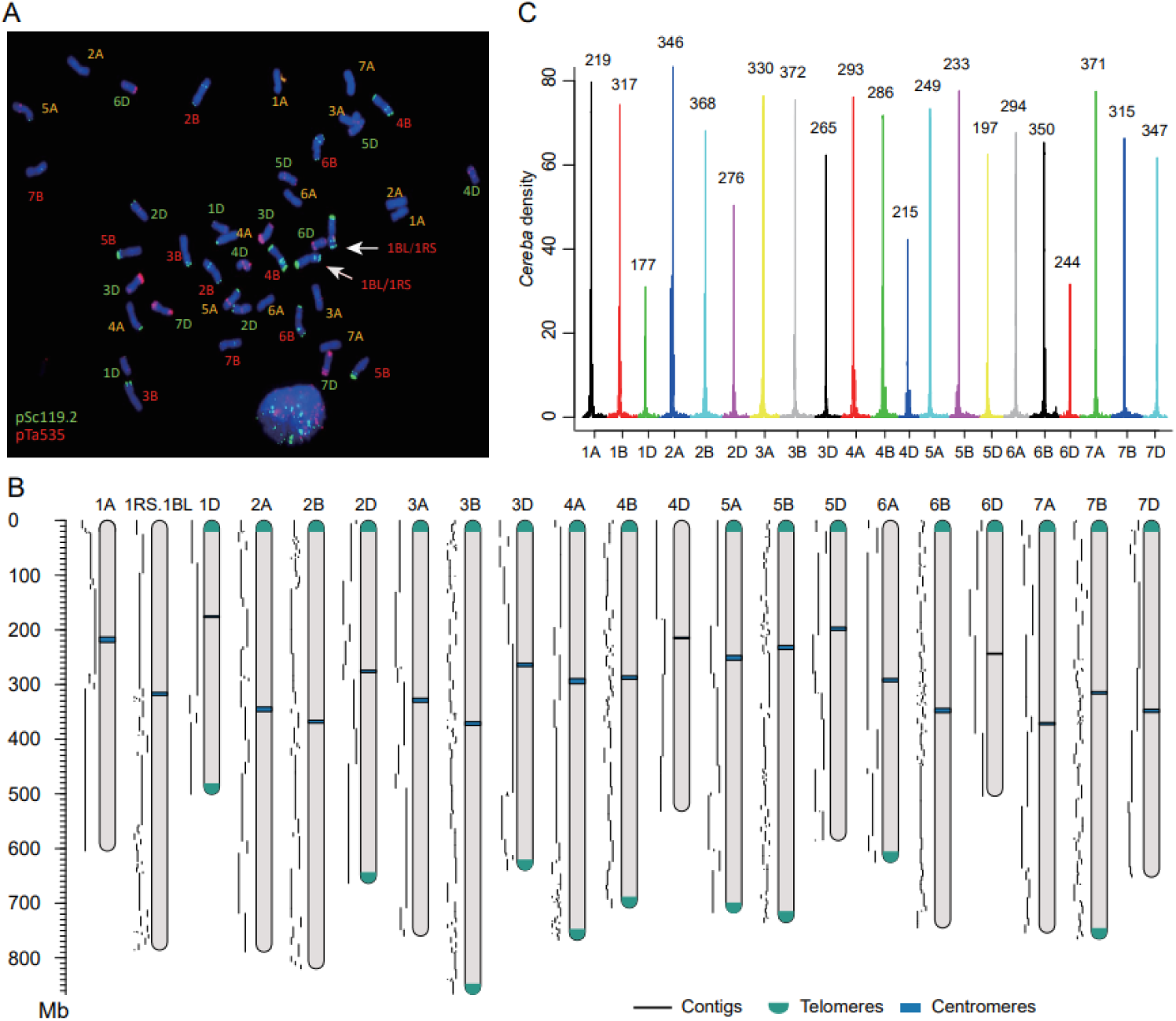
Genome assembly of Zhou8425B. **A.** Characterization of Zhou8425B somatic chromosomes by fluorescent *in situ* hybridization assays using the oligonucleotide probes pSc119.2 (green) and pTa535 (red). A total of 42 chromosomes, including two 1BL/1RS translocation chromosomes (arrowed), were consistently identified. **B.** Identification of centromeres in the assembled Zhou8425B chromosomes by examining heightened enrichment of *Cereba* transposons. The approximate physical position of centromere in each chromosome (Mb) is shown on the top of the peak. **C.** Schematic representation of the 21 assembled chromosomes of Zhou8425B. The contigs scaffolding onto each chromosome are shown on the left. Centromeres are marked by blue rectangles. Telomeres, represented by green arcs, were assembled for 18 chromosomes at either one or both ends.

**Table 2.** Main features of Zhou8425B genome assembly and annotation.

| Assembly |  |
| --- | --- |
| Assembled genome size | 14.75 Gb |
| Sequence assigned to chromosomes | 14.52 Gb |
| Number of scaffolds | 1,109 |
| N50 scaffold length | 735.11 Mb |
| N90 scaffold length | 532.03 Mb |
| Number of contigs | 1,729 |
| N50 contig length | 70.49 Mb |
| N90 contig length | 9.87 Mb |
| Longest contig | 323.62 Mb |
| Longest scaffold | 866.06 Mb |
| GC content | 46.06% |
| Annotation |  |
| Number of high-confidence genes | 106,940 |
| Mean length of high-confidence genes | 3,569 bp |
| Total length of high-confidence genes | 381.70 Mb |
| Number of genes supported by transcriptional evidence | 96,728 |
| Retrotransposons | 9.60 Gb |
| DNA transposons | 2.77 Gb |
| Other repeats | 544.37 Mb |
| Total repeat sequences | 12.91 Gb |
| Total repeat sequences percentage | 87.53% |
| Number of telomeres | 27 |
| BUSCO completeness score | 99.4% |

The assembled size of A, B, and D subgenomes was 5.01, 5.43, and 4.08 Gb, respectively (Table S4). In general, more contigs (40–85>) were needed to span individual subgenome B chromosomes, which were reduced to 11 - 42 for subgenome A chromosomes, and 5 - 15 for subgenome D chromosomes (Fig. 2B; Table S4). The largest (866.06 Mb) and smallest (500.93 Mb) assembled chromosomes were 3B and 1D, respectively (Table S4). The centromeric region, each covered by a single contig and highly enriched for *Cereba* retrotransposons (Fig. 2C), was identified for all 21 chromosomes (Table S5). A total of 27 telomeres were assembled, with 10 chromosomes (1D, 2D, 3B, 3D, 4A, 4B, 5A, 5B, 6A and 7B) having telomeres at both ends (Fig. 2B). The GC content of the assembled genome was 46.06% (Table 2).

To aid gene annotation, 12 Zhou8425B plant samples collected at the seedling, heading, and post anthesis stages were subjected to RNA sequencing, resulting in 161.35 Gb high-quality transcriptome data (Table S6). By combining these data and the gene sets reported for previously sequenced wheat genotypes, a total of 106,940 high-confidence (HC) protein-coding genes were annotated for Zhou8425B (Table 2). Nearly 99% of the 106,940 HC genes were functionally annotated by BLAST analysis in the five gene function annotation databases including NR, SwissProt, GO, KEGG, and eggnog (Table S7). The HC genes were mainly distributed towards the terminal regions of 21 chromosomes (Fig. 3). Moreover, five types of non-coding RNA genes, i.e., rRNA, tRNA, miRNA, snRNA, and snoRNA, were identified in Zhou8425B genome assembly, which amounted to 26,237, 51,883, 39,521, 1,147, and 3,973, respectively (Table S8).

**Figure 3.**
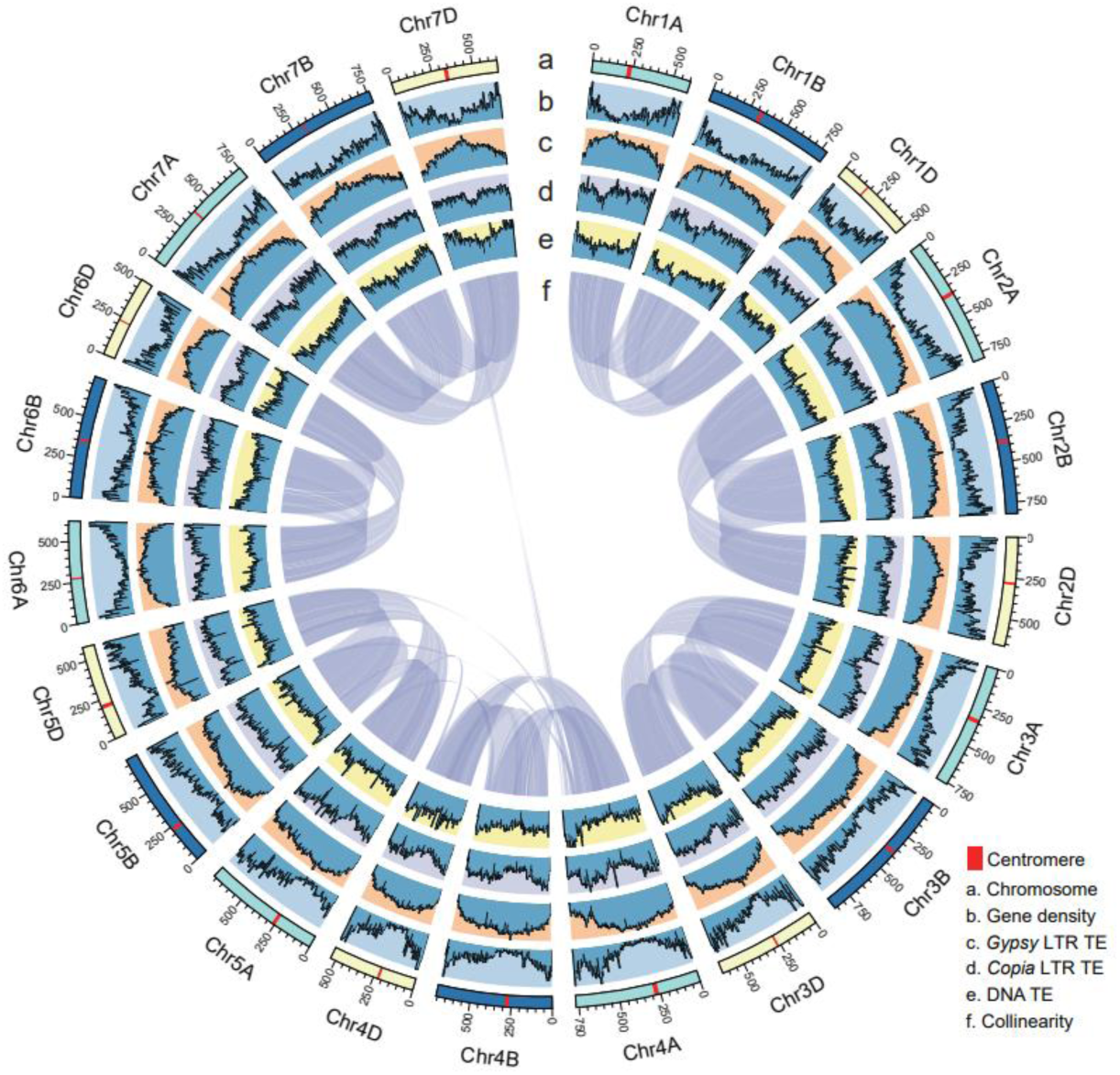
Circos plot displaying some important features of Zhou8425B genome. From the outside to the inside are the characteristics of 21 chromosomes (a), density of high confidence protein-coding genes (b), *Gypsy* LTR transposable elements (TEs) (c), *Copia* LTR TEs (d) and DNA TEs (e), and syntenic relationships in between the chromosomes of three subgenomes (f). The gene and TE densities shown in (b) to (e) were obtained by scanning the chromosome sequence in 3 Mb windows.

The total size of repetitive elements in Zhou8425B assembly was 12.91 Gb, which accounted for 87.53% of the assembled genome (Table 2; Table S9). The content of retrotransposons (9.60 Gb) was substantially larger that of DNA transposons (2.77 Gb) (Table S9). The three major families of transposons were *Gypsy*, *Copia*, and *CACTA*, which occupied 43.69%, 16.01%, and 15.05% of Zhou8425B genome, respectively (Table S9). *Gypsy* transposons tended to have higher abundance towards the centromeric region, whereas *Copia* and *CACTA* elements appeared to accumulate more in the terminal regions of chromosome (Fig. 3).

The high quality of Zhou8425B genome assembly was supported by the following lines of evidence. First, 99.96% of 43,318,145 unique HiFi reads could be properly mapped to the assembly (Table S10). Second, mapping with 368.15 Gb Zhou8425B Illumina reads (∼25X of the assembled genome) yielded a mean per base quality (QV) of 47 and an average nucleotide accuracy higher than 99.99% (Table S11). Third, a BUSCO completeness score of 99.4% was obtained for the assembly (Table S12). Finally, the long terminal repeat assembly index (LAI), which indicates the contiguity of intergenic and repetitive regions of genome assemblies (*33*), was above 16 for the A, B, and D subgenomes of Zhou8425B assembly (Fig. S2), which is higher than that reported for recently assembled hexaploid wheat genomes. Together, these data suggest that Zhou8425B genome assembly is highly contiguous and accurate in both genic and non-genic space.

To verify the high continuity and accuracy of Zhou8325B genome assembly, we analyzed the gluten gene loci, which specify two types of high-molecular-weight glutenin subunits (HMW-GSs), three types of low-molecular-weight glutenin subunits (LMW-GSs), and four types of gliadins (*34*). Variations in the amount and composition of these gluten proteins have major influences on wheat end-use properties (*34*). The gluten gene loci are highly complex, particularly those encoding LMW-GSs and/or gliadins, and are thus difficult to assemble (*34*). As anticipated, two paralogous genes specifying x- and y-types of HMW-GSs were detected in each of the three homoeologous *Glu-A1*, *-B1*, and *-D1* loci, with *Glu-A1y* being a pseudogene (Fig. S3; Table S13). Consequently, five HMW-GSs were expressed in Zhou8425B grains (Fig. S4A). In the composite locus *Gli-A1*/*Glu-A3*, 10 gluten genes including four pseudogenes were found, with the six active members coding for one δ-gliadin, two γ-gliadin, or three LMW-GS proteins (Fig. S3; Table S13). Zhou8425B lacked the *Gli-B1*/*Glu-B3* locus because of 1RS translocation that replaced the 1BS chromosome arm. We thus examined the *Sec-1* and *Sec-4* loci normally carried by 1RS (*28*). In the *Sec-1* locus, 23 gluten genes including 13 pseudogenes were present, with the 10 active members encoding 40K γ-secalins; in the *Sec-4* locus, 18 gluten genes with 15 active members encoding ω-secalins were detected (Fig. S3; Table S13). In the *Gli-D1*/*Glu-D3* locus, 17 gluten genes including four pseudogenes were observed, with the 13 active members encoding one δ-gliadin, three γ-gliadin, three ω-gliadin protein, or six LMW-GS proteins (Fig. S3; Table S13). The homoeologous *Gli-A2*, *-B2*, and *-D2* loci, resided on group 6 chromosomes, carried 15, 14, and 11 active genes encoding α-gliadins, respectively; these loci also harbored varying numbers of inactive α-gliadin genes (3–19) (Fig. S3; Table S13). Corresponding to the annotation of nine active LMW-GS genes for Zhou8425B, multiple LMW-GS proteins were found accumulated in its grains (Fig. S4B). Lastly, although the 10 gluten gene loci varied greatly in size (0.06 Mb - 20.23 Mb, Fig. S3), they were mostly covered by a single contig in Zhou8425B genome assembly, except for *Glu-B1* that was contained in two adjacent contigs (Table S13).

### Gene expression features of Zhou8425B genome assembly

Of the 106,940 HC genes, 96,728 (90.45%) was supported by transcriptional evidence (Table 2). Principal component analysis of gene expression data showed separation of 12 transcriptome samples and clustering of those with similar gene expression profiles (Fig. 4A). Based on FPKM (fragments per kilobase of exon model per million mapped fragments) values, the proportion of the genes that were expressed at low (FPKM ≤ 1), intermediate (1< FPKM ≤ 10), or high (10 < FPKM ≤ 100) level was 61.60%, 28.18%%, and 9.53%, respectively; only 0.68% of the genes were strongly expressed (FPKM > 100) (Fig. 4B). Construction of gene co-expression networks with 64,692 genes that were expressed in at least six of the 12 transcriptome sequencing samples revealed 22 distinct co-expression modules, with one to four clusters of co-expressed genes significantly associated with individual samples (Fig. 4C). More detailed analysis of the MEblue module revealed one cluster of co-expressed genes significantly associated with developing grains at 15 days post anthesis (DPA); within this cluster, a group of co-expressed genes was found with TraesZ8425B1RS01G036200, which resided on 1RS translocation and encoded a putative B3 TF, as a putative hub (Fig. 4D). The co-expressed partners of TraesZ8425B1RS01G036200 included not only the genes located on 1RS but also those on wheat chromosomes, many of which were annotated to encode gluten proteins (e.g., HMW-GSs) or the proteins involved in starch metabolism (alpha amylase inhibitor) or lipid utilization (lipid transfer proteins) (Fig. 4D; Table S14).

**Figure 4.**
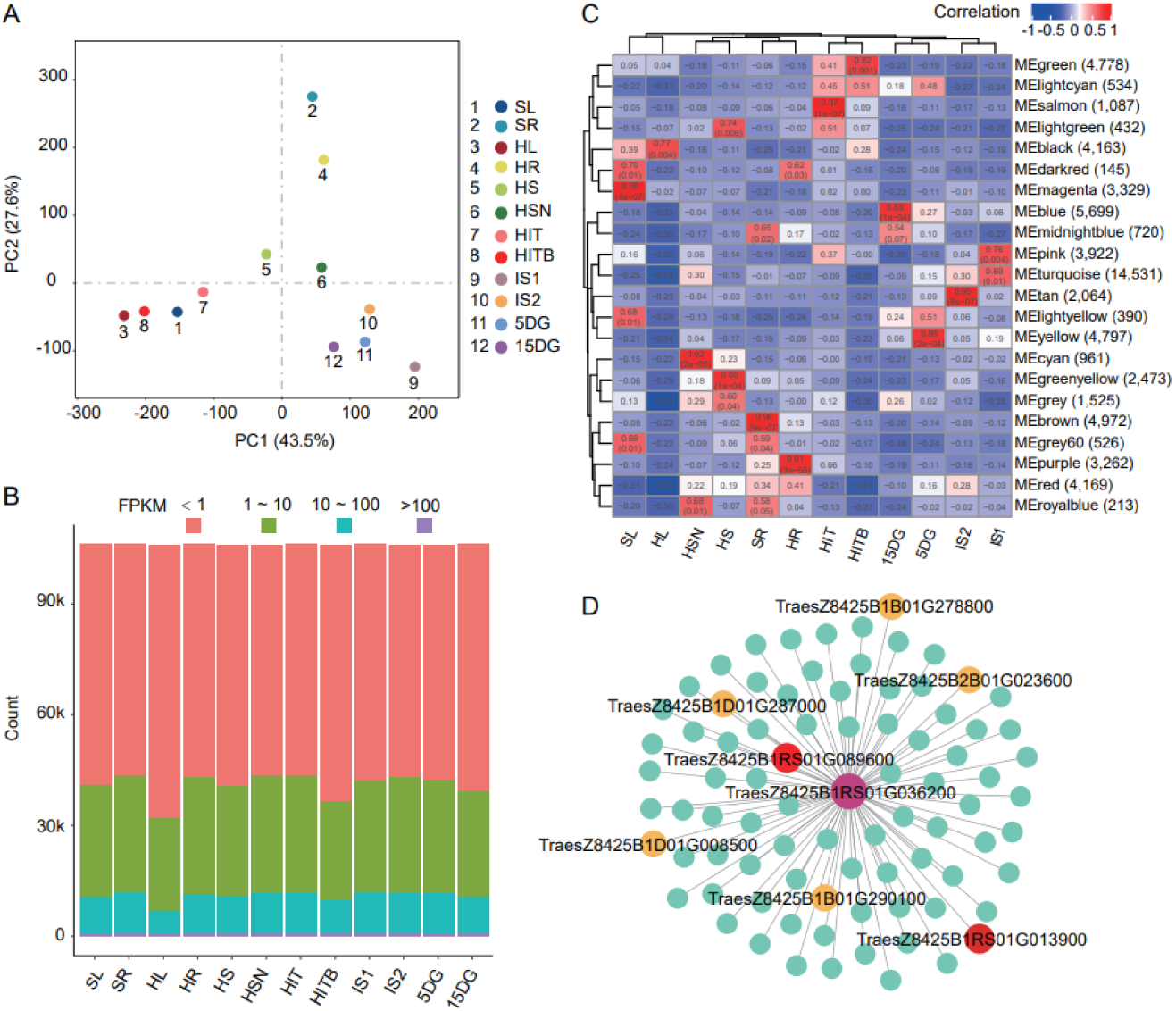
Transcriptome analysis of 12 samples collected from Zhou8425B plants. **A.** Separation and clustering of 12 samples by principal component analysis of gene expression. **B.** Differences in the expression levels of Zhou8425B genes revealed by transcriptome analysis of the 12 samples. The majority of the genes had low expression level (FPKM < 1), followed by those with intermediate (FPKM, 1 ∼ 10) or relatively high expression (FPKM, 10 ∼ 100) level. Only a minority of the genes were strongly expressed (FPKM >100). **C.** Construction of co-expression modules with the transcriptome data of 12 Zhou8425 plant samples using the R package of WGCNA. A total of 22 modules (listed on the right side) were obtained for the 12 samples (bottom panel). Each significant cell had two values, correlation coefficient (top) and *P* value (below). **D.** A subset of the genes co-expressed with TraesZ8425B1RS01G036200, which was annotated to encode a B3 transcription factor. The co-expressed partners of TraesZ8425B1RS01G036200 included those located on 1RS (e.g., TraesZ8425B1RS01G089600) or wheat chromosomes (such as TraesZ8425B1D01G287000). The seven illustrated co-expressed genes were predicted to encode high molecular weight glutenin subunit (TraesZ8425B1B01G290100), low molecular weight glutenin subunit (TraesZ8425B1D01G008500), alpha amylase inhibitor (TraesZ8425B2B01G023600), two lipid transfer proteins (TraesZ8425B1B01G278800 and TraesZ8425B1D01G287000), starch synthase (TraesZ8425B1RS01G089600), or γ-secalin (TraesZ8425B1RS01G013900). This example is extracted from the MEblue module significantly associated with developing grains at 15 days post anthesis. The 12 samples were composed of 10-day seedling leaves (SL), 10-day seedling roots (SR), heading-stage leaves (HL), heading-stage roots (HR), heading-stage stems (HS), heading-stage stem nodes (HSN), heading-stage ineffective tillers (HIT), heading-stage ineffective tiller buds (HITB), immature spikes (1 cm) (IS1), immature spikes (2 - 3 cm) (IS2), or developing grains at 5 (5DG) or 15 (15DG) days post anthesis.

### Chromosomal structural changes and tandemly duplicated gene clusters in Zhou8425B

Following the above analysis, a comparison with 16 previously assembled hexaploid wheat genomes was conducted in order to reveal unique chromosomal structural changes and highly tandemly duplicated gene clusters in Zhou8425B, which may yield important clues for studying the elite agronomic traits of Zhou8425B. The 16 varieties were originated from different parts of the world, two of which (e.g., Kenong 9204 and Aikang 58) carried 1RS translocation (Table 3). Here we focused on comparing chromosome translocations (CTs) between Zhou8428B and the 16 previously sequenced cultivars. We divided the translocations into three categories based on sizes of the translocated segments, CT ≤ 10 Mb, 10 Mb ≤ CT ≤ 50Mb, and CT ≥ 50 Mb, As shown in Table 3, the translocations were mainly smaller than 10 Mb, followed by those bigger than 10 Mb but less than 50 Mb, with the CTs larger than 50 Mb not found in most comparisons. On this basis, 27 CTs occurred in Zhou8425B but not the 16 compared lines were uncovered (Table 4).

**Table 3.** Chromosomal translocations between Zhou8425B and previously sequenced hexaploid wheat varieties.

| Variety | Translocation (Mb) |  |  |
| --- | --- | --- | --- |
|  | CT ≤ 10 | 10 ≤ CT ≤ 50 | CT ≥ 50 |
| Chinese Spring | 79 | 14 | 0 |
| Spelta | 72 | 8 | 2 |
| Kenong 9204 (1BL/1RS) | 61 | 14 | 0 |
| Aikang 58 (1BL/1RS) | 75 | 14 | 0 |
| ArinaLrFor | 85 | 21 | 11 |
| Jagger | 99 | 11 | 0 |
| Julius | 81 | 8 | 2 |
| Landmark | 89 | 8 | 2 |
| Longreach | 85 | 11 | 0 |
| Mace | 97 | 12 | 0 |
| Mattis | 82 | 18 | 10 |
| Stanely | 88 | 13 | 0 |
| Kariega | 97 | 17 | 0 |
| Renan | 29 | 0 | 0 |
| Norin61 | 87 | 11 | 0 |
| Fielder | 73 | 12 | 0 |

**Table 4.** Unique chromosomal translocations occurred in Zhou8425B genome.

| Translocation | Starting/ending position (bp)<br>(Fragment size, Mb) | Starting/<br>ending gene | Number of genes<br>included |
| --- | --- | --- | --- |
| 3A to 1B | 621,956,005/<br>625,962,171 (4.01) | TraesZ8425B1B01G280900/<br>TraesZ8425B1B01G283500 | 27 |
| 3A to 1D | 313,801,525/<br>314,035,592 (0.23) | TraesZ8425B1D01G206300/<br>TraesZ8425B1D01G206800 | 6 |
| 3A to 1D | 386,021,413/<br>391,802,228 (5.78) | TraesZ8425B1D01G266200/<br>TraesZ8425B1D01G271900 | 58 |
| 4A to 3B | 480,771,256/<br>482,619,289 (1.85) | TraesZ8425B3B01G268100/<br>TraesZ8425B3B01G269400 | 14 |
| 1B to 3D | 483,268,695/<br>485,578,878 (2.31) | TraesZ8425B3D01G341600/<br>TraesZ8425B3D01G344300 | 28 |
| 5A to 4A | 456,610,595/<br>461,712,804 (5.10) | TraesZ8425B4A01G157500/<br>TraesZ8425B4A01G160000 | 26 |
| 7A to 4A | 655,927,706/<br>664,819,470 (8.89) | TraesZ8425B4A01G351300/<br>TraesZ8425B4A01G358800 | 76 |
| 7A to 4A | 720,470,097/<br>727,395,052 (6.92) | TraesZ8425B4A01G410800/<br>TraesZ8425B4A01G422700 | 120 |
| 7A to 4A | 728,395,671/<br>732,020,762 (3.63) | TraesZ8425B4A01G422900/<br>TraesZ8425B4A01G429500 | 67 |
| 3B to 4B | 28,004,783/<br>29,286,026 (1.28) | TraesZ8425B4B01G034100/<br>TraesZ8425B4B01G036000 | 20 |
| 5A to 4B | 145,615,203/<br>151,882,831 (6.27) | TraesZ8425B4B01G124700/<br>TraesZ8425B4B01G128300 | 37 |
| 7A to 4B | 425,589,387/<br>425,903,852 (0.31) | TraesZ8425B4B01G189000/<br>TraesZ8425B4B01G190000 | 11 |
| 5A to 4B | 686,882,816/<br>689,705,768 (2.82) | TraesZ8425B4B01G376300/<br>TraesZ8425B4B01G380600 | 44 |
| 5A to 4B | 690,220,997/<br>690,280,792 (0.06) | TraesZ8425B4B01G380900/<br>TraesZ8425B4B01G381300 | 5 |
| 5A to 4B | 705,236,021/<br>705,583,062 (0.35) | TraesZ8425B4B01G403900/<br>TraesZ8425B4B01G405000 | 12 |
| 3B to 4D | 15,078,325/<br>16,864,616 (1.79) | TraesZ8425B4D01G029700/<br>TraesZ8425B4D01G032400 | 28 |
| 5A to 4D | 102,616,067/<br>110,224,794 (7.61) | TraesZ8425B4D01G114600/<br>TraesZ8425B4D01G118700 | 42 |
| 5A to 4D | 495,568,145/<br>523,772,560 (28.2) | TraesZ8425B4D01G311500/<br>TraesZ8425B4D01G360800 | 494 |
| 5A to 4D | 524,127,339/<br>531,789,532 (7.66) | TraesZ8425B4D01G363000/<br>TraesZ8425B4D01G380000 | 171 |
| 5A to 4D | 530,820,380/<br>531,258,448 (0.44) | TraesZ8425B4D01G378100/<br>TraesZ8425B4D01G379400 | 14 |
| 4A to 5B | 12,638,955/<br>14,801,180 (2.16) | TraesZ8425B5B01G006500/<br>TraesZ8425B5B01G008000 | 16 |
| 4A to 5B | 293,349,592/<br>299,972,064 (6.62) | TraesZ8425B5B01G144500/<br>TraesZ8425B5B01G148800 | 44 |
| 4A to 5B | 688,238,048/<br>733,019,911 (44.78) | TraesZ8425B5B01G490200/<br>TraesZ8425B5B01G554600 | 645 |
| 1A to 5B | 705,439,596/<br>708,027,603 (2.59) | TraesZ8425B5B01G515900/<br>TraesZ8425B5B01G519300 | 35 |
| 1A to 5D | 564,075,486/<br>565,524,236 (1.45) | TraesZ8425B5D01G495600/<br>TraesZ8425B5D01G497700 | 22 |
| 4A to 5D | 247,033,266/<br>252,028,777 (4.99) | TraesZ8425B5D01G135000/<br>TraesZ8425B5D01G137600 | 27 |
| 4A to 5D | 549,902,717/<br>576,012,970 (26.11) | TraesZ8425B5D01G473200/<br>TraesZ8425B5D01G516200 | 431 |

Most of these unique CTs were less than 10 Mb (0.06 - 8.89 Mb), with the genes carried by these translocated segments being less than 120 (Table 4). However, three of them were relatively large (26.11 - 44.78 Mb), with the residing genes varying from 431 to 645 (Table 4). Notably, 11 of 27 unique CTs involved transfers of chromosomal fragments from subgenomes A or B to subgenome D (Table 4). Comparison of gene expression between the translocated fragments (in Zhou8425B) and their untranslocated counterparts (in Aikang 58) showed that genes in the translocated fragments tended to be more highly expressed (Fig. 5).

**Figure 5.**
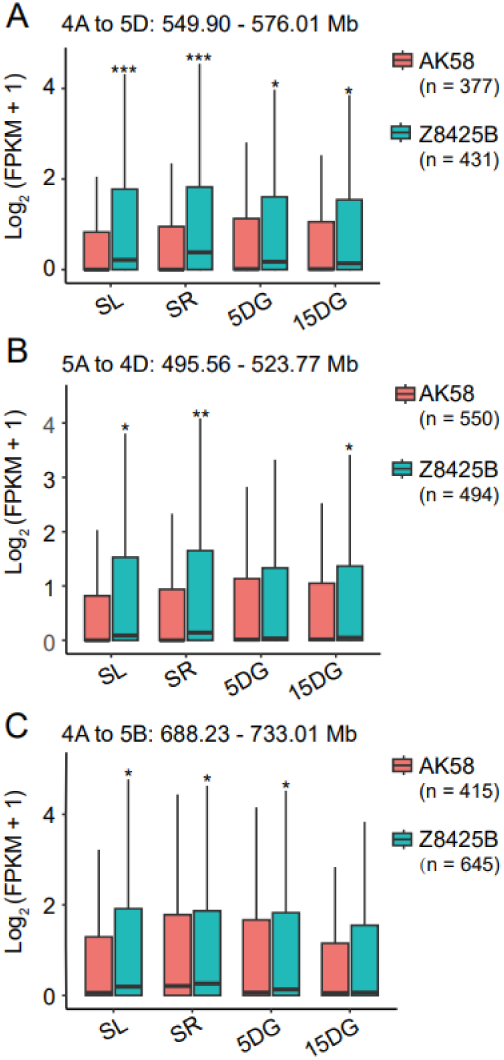
Expression divergence between the genes in the translocated segment and those in the untranslocated orthologous counterpart. The three analyzed segments translocated in Zhou8425B (Z8425B), i.e., 4A to 5D (**A**), 5A to 4D (**B**), or 4A to 5B (**C**), but not in Aikang 58 (AK58). In general, the genes in the translocated segments tended to be more highly expressed relative to those in the untranslocated orthologous counterparts based on analyzing the transcriptome data of four different plant samples. The values in the brackets indicate numbers of genes carried by the compared segments. SL, 10-day seedling leaves; SR, 10-day seedling roots, 5DG, developing grains at 5 days post anthesis (DPA); 15DG, developing grains at 15 DPA. Statistical analysis was conducted using Student *t* test. *, *P* < 0.05; **, *P* < 0.01; ***, *P* < 0.001.

Tandemly duplicated genes play crucial roles in regulating the growth, development, and environmental adaptations of high plants in response to internal and external cues (*35*). They often exist as clusters with multiple paralogous members in each cluster. Resolving genotype-dependent variations of tandemly duplicated gene clusters (TDGCs) may yield important information for studying and improving agronomic traits. We therefore computed TDGCs in Zhou8425B and 16 previously sequenced common wheat cultivars and divided the resultant TDGCs into four types, Types 1 to 4 with their paralogous members being 3 - 5, 6 - 16, 17 - 25, and 25 plus, respectively (Fig. 6A). Types 1 to 3 TDGCs were found in all 17 cultivars, with Type 4 having the highest number, followed by Type II and then Type III (Fig. 6A). In contrast, Type IV TDGCs were detected in only three cultivars (Fielder, Kariega, and Zhou8425B), and their numbers were all smaller than five (Fig. 6A). The three super Type IV TDGCs (STDGCs) of Zhou8425B were located on 2DL and 4BL, respectively (Fig. S5). Amino acid sequence analysis indicated that paralogs in the 2DL-1 and 2DL-2 STDGCs may encode cysteine proteases and flavin-binding monooxygenase-like family proteins, respectively, while those in the 4BL STDGC could specify germin-like proteins (GLPs) (Fig. 6B). Although the paralogs in 2DL-1 and 2DL-2 STDGCs did not show appreciable levels of expression in the 12 transcriptome samples sequenced in this work, those of the 4BL STDGC exhibited preferential expression in the roots at both seedling and heading stages (Fig. 6C).

**Figure 6.**
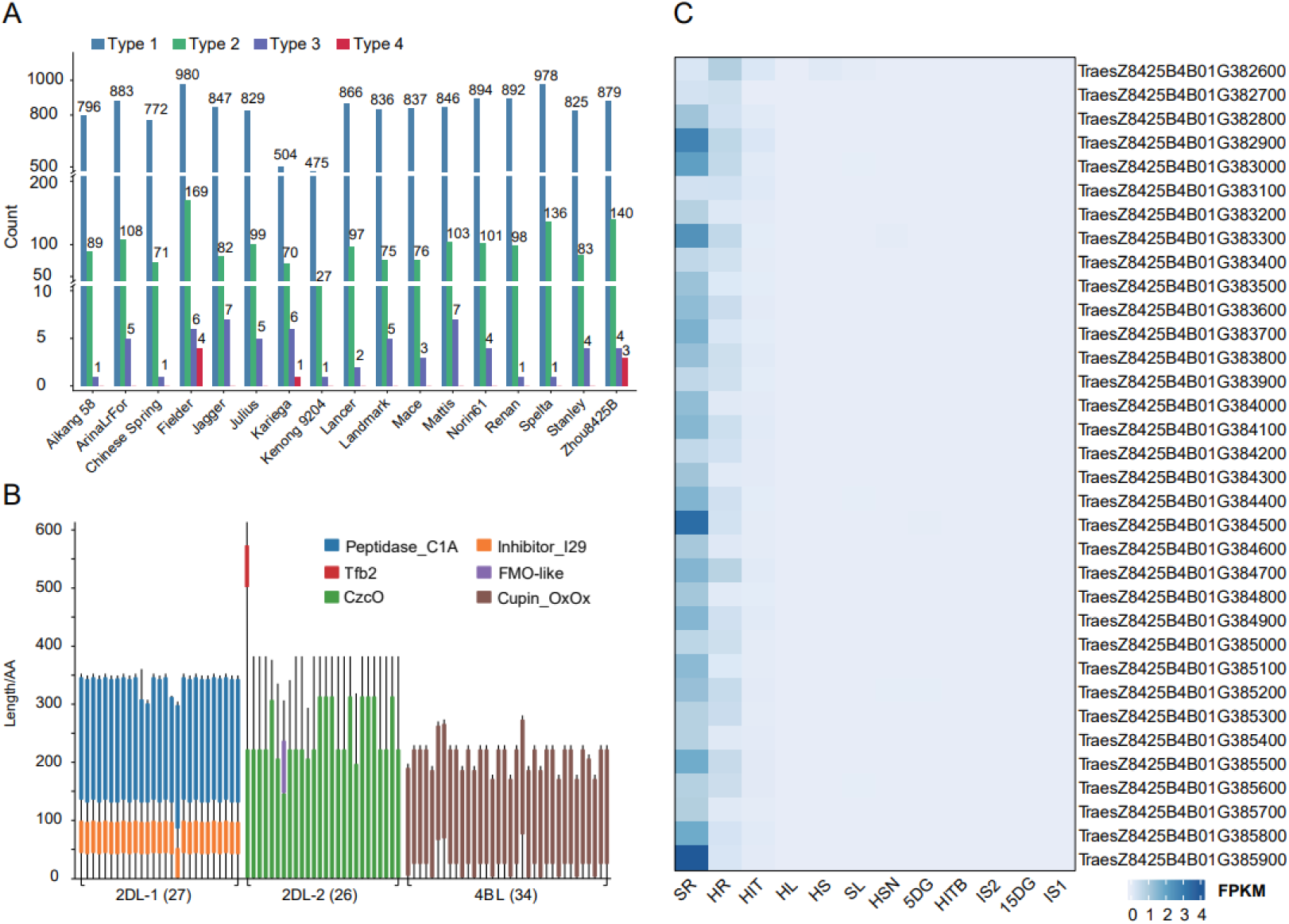
Analysis of tandemly duplicated gene clusters. **A.** Different types of tandemly duplicated gene clusters (TDGCs) detected in the genomes of Zhou8425B and 16 other common wheat varieties. Types 1 to 4 refer the TDGCs with varying numbers of constituent paralogs, i.e., Type 1 ≤ 5, 5 < Type 2 ≤ 15, 15 < Type 3 ≤ 25, and Type 4 > 25. **B.** Protein domains carried by paralogs of the three super TDGCs (2DL-1, 2DL-2, and 4BL) detected in Zhou8425B. These super TDGCs had 17, 26, and 34 paralogs, respectively. **C.** Expression heatmap of the 34 paralogs in the 4BL super TDGC prepared using the transcriptome sequencing data of 12 Zhou8425B plant samples. These paralogs showed preferential expression in wheat roots at both seedling and heading stages. The abbreviations of the 12 samples are explained in Figure 4.

### Distinct features of Zhou8425B 1RS translocation

The 1RS translocation has gained extensive use in worldwide wheat breeding since 1970s and has contributed substantially to global wheat improvement (*27, 36*). Although resistance genes (*Pm8, Yr9, Lr26,* and *Sr31*) carried by the 1RS originated from Petkus rye have lost their functions due to changes in pathogen races, there are several other sources of 1RS that have been reported to confer resistance to multiple pathogens after being transferred to wheat through distant hybridization (*36, 37*). Furthermore, there is increasing evidence that 1RS also carries the genes beneficial for wheat tolerance to abiotic stresses such as drought (*38, 39*). Despite its importance to wheat improvement, detailed genomic analysis of 1RS translocations in different common wheat backgrounds is still lacking. Recently, genome assemblies for two common wheat cultivars (Kenong 9204 and Aikang 58) carrying 1RS translocation and two diploid rye lines (Lo7 and Weining) have been reported (*8, 9, 27, 28*). We thus compared the 1RS of Zhou8425B with its counterparts in Kenong 9204, Aikang 58, Lo7, and Weining.

Pairwise comparisons revealed that the size of Zhou8425B 1RS (317 Mb) was obviously longer than that in Kenong 9204 (275 Mb), Aikang 58 (282 Mb), Lo7 (267 Mb), and Weining (310 Mb); although these 1RS arms were largely colinear, there existed complex chromosome structural changes, including inversions and translocations (Fig. 7A). Notably, a non-collinear region of approximately 20.5 Mb, located in the telomere end of Zhou8425B 1RS, was consistently found in the comparisons (Fig. 7A). Further analysis revealed that this segment was primarily composed of three types of tandemly repeated satellite arrays with the basic repeat unit being 380, 571, and 118 bp, respectively (Fig. 7B). Remarkably, this type of highly repetitive segment was not found in the previously sequenced 18 common wheat and rye genotypes.

**Figure 7.**
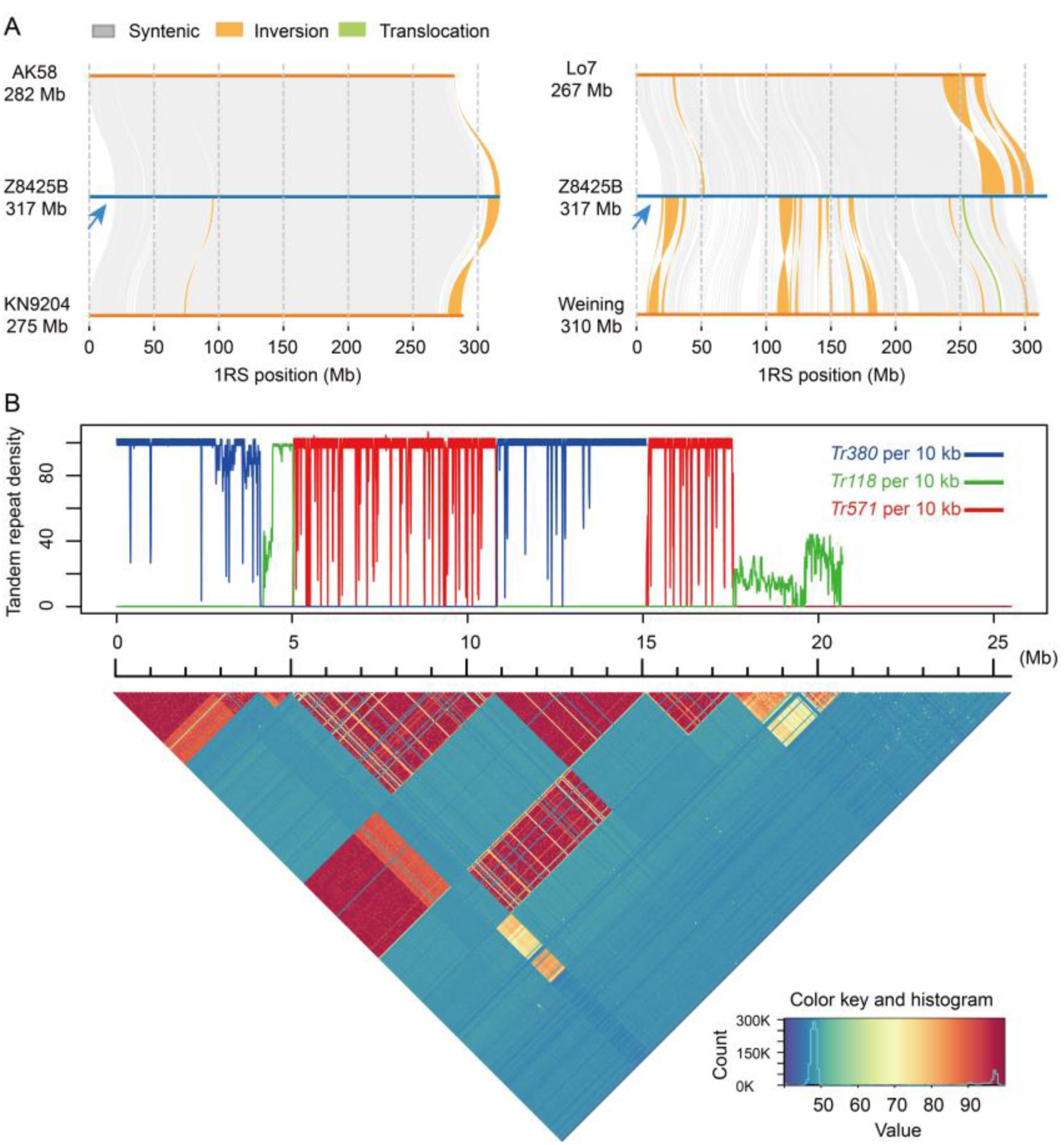
Analysis of structural variations of 1RS chromosome arm. **A.** Syntenic relationships between the 1RS in Zhou8425B and its counterparts in two common wheat cultivars, Aikang 58 (AK58) and Kenong 9204 (KN9204), and two diploid rye lines (Lo7 and Weining). The five 1RS arms differed in length, with a non-collinear segment (∼ 20.5 Mb) present in the telomere end of Zhou8425B 1RS (arrowed). **B.** Presence and arrangement of three types of tandemly repeated satellite arrays in the 20.5 Mb non-collinear segment. The heatmap below shows pairwise sequence identities between all nonoverlapping 5-kb windows in the 20.5 Mb non-collinear segment.

The numbers of annotated HC genes and computed gene families differed among the five compared 1RS arms (Fig. 8A). A total of 440 gene families were conserved among the 1RS arms, with 9 - 46 specific gene families detected for them (Fig. 8B). Interestingly, the number of genes predicted to encode B3 or AP2/ERF-ERF TFs, which frequently regulate plant tolerance to abiotic stresses (*40, 41*), were expanded in Zhou8425B relative to Kenong 9204, Aikang 58, Lo7, Weining (Fig. 8C). The *AP2/ERF-ERF* genes of Zhou8425B 1RS were also more abundant compared to homoeologous 1BS of the sequenced common wheat lines (Fig. 8C). Notably, one of the B3 genes, TraesZ8425B1RS01G036200, was more highly expressed in the developing grains at 15 DPA (Fig. 8D); this gene had a large number of co-expressed partners in developing grains (Fig. 4D). Moreover, one of the AP2/ERF-ERF genes, TraesZ8425B1RS099000, was expressed in multiple plant samples, especially those composed of immature spikes, heading stage stems, or heading stage immature tillers (Fig. 8D).

**Figure 8.**
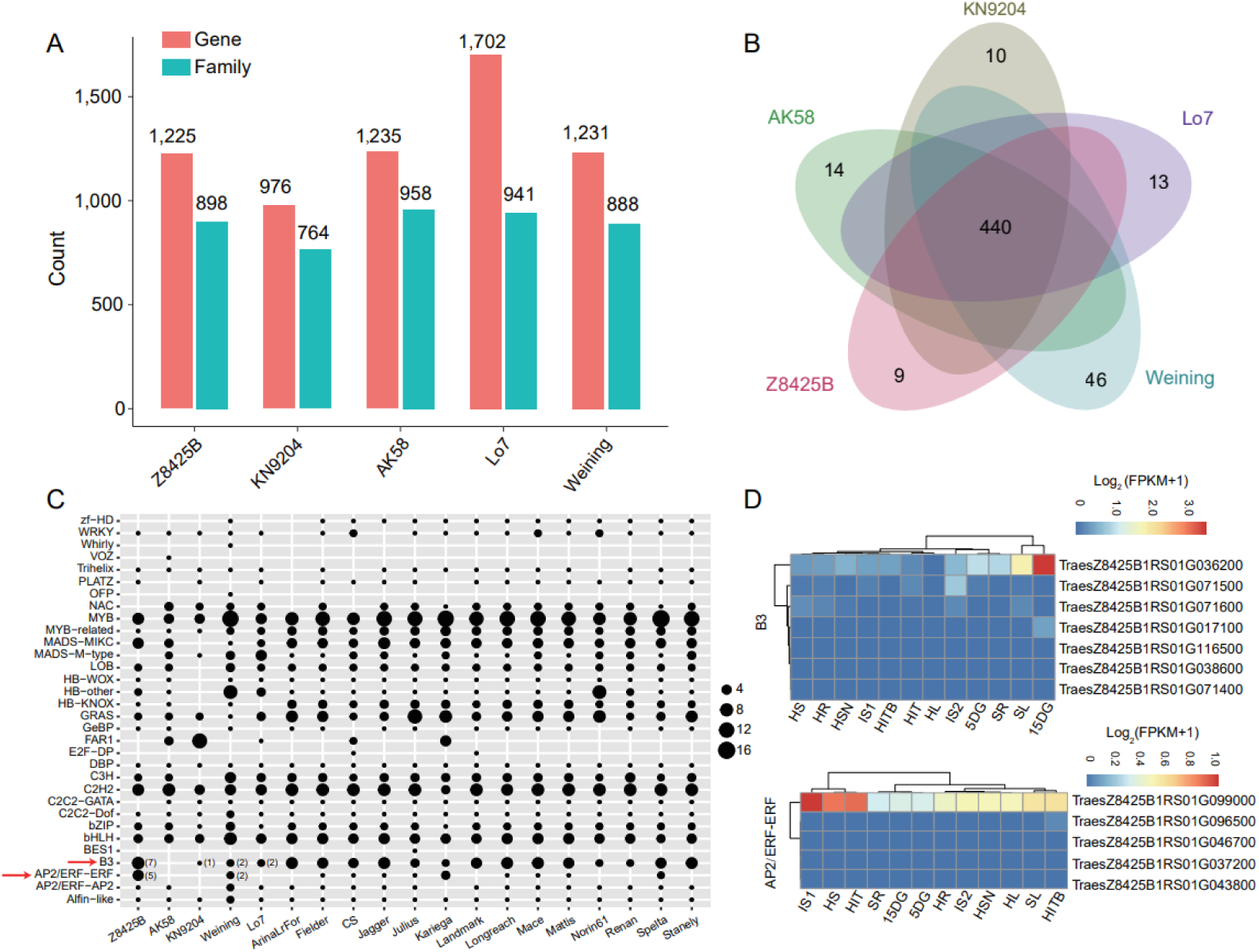
Comparison of gene contents, gene families, and transcription factor genes among five 1RS arms. **A.** The numbers of genes and gene families annotated for the 1RS arms in Zhou8425B (Z8425B), Kenong 9204 (KN9204), Aikang 58 (AK58), Lo7, and Weining. **B.** Conserved and unique gene families computed for the 1RS arms in Zhou8425B (Z8425B), Kenong 9204 (KN9204), Aikang 58 (AK58), Lo7, and Weining. **C.** Expansion of the genes encoding B3 or AP2/ERF-ERF TFs (arrowed) in the 1RS of Zhou8425B (Z8425B) relative to the 1RS in Aikang 58 (AK58), Kenong 9204 (KN9204), Weining, and Lo7. The *AP2/ERF-ERF* genes were also more numerous for Z8425B 1RS compared to homoeologous 1BS arm of previously sequenced common wheat varieties. **D.** Expression heatmaps of the genes encoding B3 or AP2/ERF-ERF TFs carried by the 1RS of Zhou8425B. The heatmaps were created with the transcriptome sequencing data of 12 Zhou8425B plant samples, whose abbreviations are explained in Figure 4.

### Accelerated mapping of YrZH3BS aided by Zhou8425B genome assembly

By combining genome-wide association study (GWAS) and Zhou8425B genome sequence assisted bi-parental mapping, we identified a new APR locus (i.e., *YrZH3BS*) conferring YR resistance and mapped it a 1 - 2 Mb interval on the short arm of chromosome 3B. The initial GWAS experiment was conducted with 245 wheat cultivars grown in five different field environments with natural and artificially induced YR infections. Flag leaf disease severity (FLDS), calculated as percentage of leaf area covered by *Pst* pustules, was scored for each line. The resultant FLDS values were used in GWAS analysis in conjunction with 612,673 SNP markers with known physical locations (Fig. S6). A total of non-redundant 214 SNPs significantly associated FLDS were detected using the phenotypic data collected from five environments with two biological repeats in each, which were mainly distributed in several GWAS peaks resided on 3BS, 3DS, 7BL, or 7DL (Tables S15 and S16).

We then focused on the significantly associated SNPs located in the interval of 6.95 - 21.40 Mb of 3BS (Fig. 9A). Haplotype analysis showed that the 14 associated SNPs in this block formed three major haplotypes, with the mean FLDS value of Hap C (19.19) being significantly (*P* < 0.0001) lower than those of Hap A (50.25) and Hap B (55.40) (Fig. 9B). These data pointed to the presence of a new APR locus on 3BS (*YrZH3BS*). To verify *YrZH3BS*, further bi-parental mapping with 198 F_2:3_ lines, developed using Xinhuamai 818 (Hap C) and Yumai 1 (Hap A) as parents, was carried out. Xinhuamai 818, derived from Zhou8425B (Fig. 9C), was highly resistant (mean FLDS = 4.50%) to YR, whereas Yumai 1 was strongly susceptible (mean FLDS = 84.17%) in the field (Fig. 9D). Based on FLDS values, the 198 F_2:3_ lines could be divided into three groups, 42 homozygously resistant, 101 heterozygously resistant, and 56 homozygously susceptible (Table S17), which fitted a 1:2:1 segregation ratio (χ^2^ = 2.02 < χ^2^_0.05, 2_ = 5.99). This validated the existence of *YrZH3BS* as a functional APR gene against YR.

**Figure 9.**
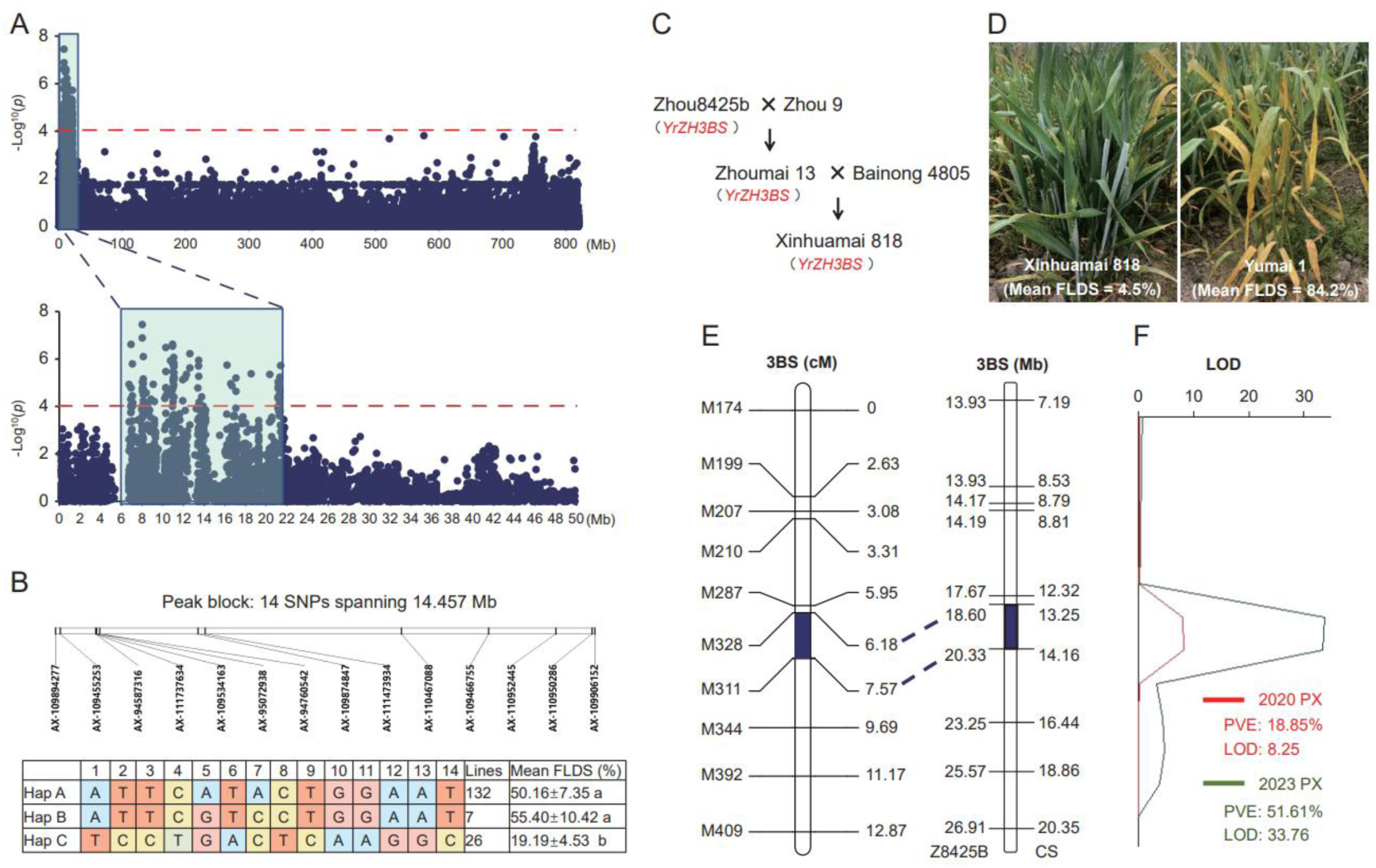
Discovery and fine mapping of a new APR locus (*YrZH3BS*) against wheat yellow rust disease. **A.** Manhattan plot showing a GWAS peak significantly associated with flag leaf disease severity (FLDS). This peak was located on 3BS chromosome arm. **B.** Differences in mean FLDS values among three major haplotypes formed by the 14 SNPs located in the 3BS GWAS peak. FLDS values were means ± SEs, which were statistically compared by Duncan’s test after analysis of variance, with significant difference indicated by dissimilar letters after the FLDS values. **C.** Pedigree of Xinhuamai 818. The *YrZH3BS* locus in Xinhuamai 818 was inherited from Zhou8425B. **D.** Comparison of yellow rust resistance and FLDS between Xinhuamai 818 and Yumai 1. Xinhuamai 818, but not Yumai 1, was highly resistant to yellow rust in the field. Consistently, the mean FLDS value of Xinhuamai 818 was substantially lower than that of Yumai 1. **E.** Fine mapping of *YrZH3BS* to a 1.39 cM interval on 3BS using 10 polymorphic DNA markers developed with the aid of Zhou8425B genome sequence. The 1.39 cM interval corresponds to a physical distance of 1.73 Mb in Zhou8425B 3BS and 0.91 Mb in CS 3BS. This mapping was conducted with the F_2:3_ lines prepared using Xinhuamai 818 and Yumai 1 as parents. **F.** The percentages of FLDS variance explained by *YrZH3BS* in two field environments (2020PX and 2023PX). The *YrZH3BS* QTLs detected in the two environments were highly significant based on their respective LOD values.

To finely map *YrZH3BS*, we made use of the genome sequence information of Zhou8425B and Chinese Spring (CS) to accelerate the development of polymorphic DNA markers. Zhou8425B was especially valuable for this purpose, because it not only possessed durable APR to YR but also served as a parent for breeding Zhoumai 13 that was used for developing Xinhuamai 818 (Fig. 9C). The 3BS sequence in Zhou8425B and CS was analyzed, with the resultant indels exploited to develop DNA markers. After mapping in Xinhuamai 818 and Yumai 1 and derivative F_2:3_ lines, 10 polymorphic markers (M174, M199, M207, M287, M328, M311, M344, M392, and M409, Table S18) were genetically assigned to the initial genomic interval associated with *YrZH3BS*, with M328 and M311 being the closest flanking markers (Fig. 9E). The genetic distance covered by the two markers was 1.39 cM, which corresponds to a physical distance of 1.73 Mb in Zhou8425B and 0.91 Mb in CS (Fig. 9E). In this bi-parental mapping analysis, *YrZH3BS* explained 18.85% - 51.60% of the variance of FLDS in two different field environments (Fig. 9F).

### Individual and combined effects of three APR loci in decreasing YR disease severity and prompting grain yield traits

The finding of *YrZH3BS* raised the question if there might be positive interactions among *YrZH3BS*, *YrZH22*, and *YrZH84* in decreasing YR disease severity and promoting grain yield traits. To investigate this question, we collected 212 common wheat cultivars developed by using Zhou8425B or its early descendants (Zhoumai 13, Zhoumai 16, Zhoumai 22, and Aikang 58) as parents, and grew them in two field environments with natural and artificially induced YR infections. FLDS and grain yield related traits, including spike number per plant (SNPP), grain number per spike (GNPS), thousand grain weight (TGW), and grain yield per m^2^ (GY), were scored for each line in each environment (Table S19). Concomitantly, the 212 cultivars were genotyped for the presence of *YrZH3BS*, *YrZH22*, and/or *YrZH84* using their respective molecular markers (Table S20), which identified the cultivars lacking all three genes or carrying one, two or all three genes (Table S19).

Joint analysis of genotypic and phenotypic data revealed that presence of *YrZH3BS* (n = 84), *YrZH22* (n = 140) or *YrZH84* (n = 72) significantly decreased FLDS (by 31.37% - 55.10%) and enhanced GNPS (6.20% - 8.30%), TGW (12.00% - 17.31%), and GY (14.88 - 22.77%) (Table 5). The three genes formed three types of double combinations, with *YrZH3BS*/*YrZH22*, *YrZH3BS*/*YrZH84*, *YrZH22*/*YrZH84* detected in 66, 35, and 43 cultivars, respectively, which decreased FLDS by 52.08% - 71.11% and enhanced GNPS (7.59% - 8.49%), TGW (18.23% - 24.75%), and GY (21.76 - 29.62%) (Table 6). Triple combination of the three genes (*YrZH3BS*/*YrZH22*/*YrZH84*) was detected in only 26 cultivars, which strongly decreased FLDS (by 84.09%) and substantially elevated GNPS (8.29%), TGW (29.76%), and GY (31.98%) (Table 6). Hence, the triple gene combination appeared to be most effective in lowering FLDS and raising GNPS, TGW, and GY, followed by the double gene combinations and then the single genes (Tables 5 and 6). Finally, either single or combinations of the three genes failed to affect SNPP, although it consistently benefited GNPS, TGW, and GY (Tables 5 and 6).

**Table 5.** Effects of three APR loci on decreasing FLDS and enhancing yield related traits.

|  | Genotype |  |  |  |  |  |
| --- | --- | --- | --- | --- | --- | --- |
|  | -YrZH3BS<br>(n = 128) | +YrZH3BS<br>(n = 84) | -YrZH22<br>(n = 72) | +YrZH22<br>(n = 140) | -YrZH84<br>(n = 140) | +YrZH84<br>(n = 72) |
| FLDS | 0.48 ± 0.02 a | 0.28 ± 0.03 b<br>(-41.67%) | 0.51 ± 0.03 a | 0.35 ± 0.02 b<br>(-31.37%) | 0.49 ± 0.02 a | 0.22 ± 0.03 b<br>(-55.10%) |
| SNPP | 7.54 ± 0.02 a | 7.53 ± 0.02 a | 7.57 ± 0.03 a | 7.52 ± 0.02 a | 7.53 ± 0.02 a | 7.55 ± 0.02 a |
| GNPS | 35.79 ± 0.37 b | 38.01 ± 0.51 a<br>(+6.20%) | 35.37 ± 0.54 b | 37.33 ± 0.37 a<br>(+5.54%) | 35.66 ± 0.38 b | 38.62 ± 0.44 a<br>(+8.30%) |
| TGW<br>(g) | 35.81 ± 0.49 b | 40.97 ± 0.81 a<br>(+14.41%) | 35.07 ± 0.67 b | 39.28 ± 0.59 a<br>(+12.00%) | 35.75 ± 0.51 b | 41.94 ± 0.77 a<br>(+17.31%) |
| GY<br>(g/m<sup>2</sup>) | 382.73 ± 8.01 b | 448.38 ± 12.11 a<br>(+17.15%) | 372.17 ± 10.08 b | 427.54 ± 9.12 a<br>(+14.88%) | 379.40 ± 8.49 b | 465.79 ± 10.16 a<br>(+22.77%) |

Values were calculated as means ± SE. Statistical analysis was performed using Student *t* test. FLDS, flag leaf disease severity; SNPP, spike number per plant; GNPS, grain number per spike; TGW, thousand grain weight; GY, grain yield. FLDS was calculated as the percentage of flag leaf area with yellow rust pustules.

**Table 6.** Effects of double and triple combinations of three APR loci on decreasing FLDS and enhancing yield related traits.

| Genotype |  |  |  |  |  |  |  |  |
| --- | --- | --- | --- | --- | --- | --- | --- | --- |
|  | *-YrZH3BS-YrZH22*<br>(n = 146) | *+YrZH3BS+YrZH22*<br>(n = 66) | *-YrZH3BS-YrZH84*<br>(n = 177) | *+YrZH3BS+YrZH84*<br>(n = 35) | *-YrZH22-YrZH84*<br>(n = 169) | *+YrZH22+YrZH84*<br>(n = 43) | *-YrZH3BS-YrZH22-YrZH84*<br>(n = 128) | *+YrZH3BS+YrZH22+YrZH84*<br>(n = 26) |
| FLDS | 0.48±0.02 a | 0.23±0.03 b<br>(-52.08%) | 0.45±0.02 a | 0.13±0.04 b<br>(-71.11%) | 0.46±0.02 a | 0.14±0.03 b<br>(-69.57%) | 0.44±0.02 a | 0.07±0.03 b<br>(-84.09%) |
| SNPP | 7.54±0.02 a | 7.52±0.02 a | 7.54±0.02 a | 7.52±0.03 a | 7.53±0.02 a | 7.54±0.03 a | 7.54±0.02 a | 7.52±0.03 a |
| GNPS | 35.82±0.37 b | 38.54±0.49 a<br>(+7.59%) | 36.18±0.41 b | 39.13±0.69 a<br>(+8.15%) | 36.05±0.36 b | 39.11±0.43 a<br>(+8.49%) | 36.30±0.41 b | 39.31±0.49 a<br>(+8.29%) |
| TGW (g) | 35.82±0.48 b | 42.35±0.85 a<br>(+18.23%) | 36.44±0.55 b | 45.01±1.21 a<br>(+23.52%) | 36.04±0.45 b | 44.96±0.88 a<br>(+24.75%) | 36.52±0.53 b | 47.39±1.08 a<br>(+29.76%) |
| GY (g/m<sup>2</sup>) | 382.81±7.68 b | 466.11±12.95 a<br>(+21.76%) | 392.17±9.14 b | 492.53±15.06 a<br>(+25.59%) | 385.57±7.56 b | 499.79±10.94 a<br>(+29.62%) | 393.31±8.74 b | 519.09±13.18 a<br>(+31.98%) |

Values were calculated as means ± SE. Statistical analysis was performed using Student *t* test. FLDS, flag leaf disease severity; SNPP, spike number per plant; GNPS, grain number per spike; TGW, thousand grain weight; GY, grain yield. FLDS was calculated as the percentage of flag leaf area with yellow rust pustules.

## Discussion

Genetic improvement of crop plants has entered the era of genomics-assisted breeding (*42, 43*). High-quality genome sequence information is an essential prerequisite for efficiently deciphering and improving crop traits. Sequencing elite wheat cultivars has directly aided the mining of many important genes and loci controlling disease resistance, nutrient use efficiency, and yield related traits (*6–9*). In this study, we constructed a highly contiguous and accurate genome assembly for Zhou8425B, a key wheat breeding germplasm with many hundreds of derivative commercial cultivars (Table S1). Consistent with its superior agronomic traits (Fig. 1), Zhou8425B carries multiple elite genes and QTL alleles, including those regulating plant height, photoperiod insensitivity, APR against YR and LR diseases, and yield related traits as revealed in this work (Table 1) and previous studies.

Based on contig and scaffold N50 (Table 2), BUSCO completeness score (Table S12), and LAI values (Fig. S2), the quality of Zhou8425B genome assembly is substantially improved relative to that of previously reported common wheat genomes (*3–9*), with each chromosome covered by 5 - 85 contigs (Fig. 2C; Table S4). This is mainly due to integrated use of powerful HiFi long reads and effective Hi-C sequencing data, a strategy that has been found efficient for developing highly contiguous polyploid genome assemblies (*44*). Owing to the large genome size of hexaploid wheat and prohibitive sequencing costs, we did not use Nanopore sequencing technology in this work, thus missing the opportunity to obtain a telomere-to-telomere genome assembly at present. Nevertheless, the current version of Zhou8425B assembly could resolve highly complex genomic regions, such as centromeres, large and complex gluten gene loci, and STDGCs.

According to the transcriptome data generated in this work, more than 90% of the HC protein-coding genes annotated for Zhou8425B are expressed, although their expression levels differ considerably in different organs at different developmental stages (Fig. 4A, B). We constructed 22 co-expression modules, and illustrated their usefulness for investigating gene expression networks by analyzing a cluster of genes co-expressed in the developing grains with TraesZ8425B1RS01G036200 as a putative hub (Fig. 4C, D). Although resided on 1RS, the co-expressed partners of TraesZ8425B1RS01G036200 included not only the genes located on 1RS but also those on wheat chromosomes (Fig. 4D). Previous studies have shown that translocated rye genes function in wheat root growth and drought tolerance (*38, 39*). This encourages us to invest more effort to confirm the hub function of TraesZ8425B1RS01G036200 in further research. The result may shed new light on the contributions of translocated rye genes to the genetic control of wheat agronomic traits.

Segmental translocations are an important type of chromosomal structural changes that have been found to affect biological traits by altering gene expression (*45*). Through comparative analysis of genome assemblies, we show that extensive CTs, especially those smaller than 10 Mb, occur between different wheat cultivars (Table 3). Genotype specific CTs, identified through this type of analysis, are fairly numerous (27 for Zhou8425B, Table 4). Remarkably, the genes in the translocated segments tended to be more highly expressed relative to those in the untranslocated orthologous regions (Fig. 5). Together, these findings may stimulate more systematic studies of the roles of CTs in shaping wheat agronomic traits in the future. We noted that many of the private CTs of Zhou8425B involved transfers of chromatins from subgenomes A/B to D subgenome (Table 4). This is surprising considering that the hybridization between free-threshing tetraploid wheat and the D genome donor *Ae. tauschii*, which gave rise to hexaploid wheat, happened only around 10,000 years ago. One possible explanation is that a gamma ray irradiation treatment was used during the breeding of Zhou8425B (Fig. 1A), which might promote such CTs. However, further work is needed to verify this possibility.

Tandem gene duplication can lead to production and expression of multiple paralogs, which is an important way to enhance the strength and diversity of gene function (*35*). Our analysis suggests that wheat cultivars vary substantially in accumulating different types of TDGCs, with STDGCs (> 25 paralogs in each cluster) occurred in relatively few cultivars including Zhou8425B (Fig. 6A). Interestingly, the 4BL STDGC of Zhou8425B, with 34 paralogs annotated to encode GLPs, are preferentially expressed in the roots (Fig. 6B, C). In line with our finding, a previous study also observed strong tandem duplication of *GLP* genes on the 4BL chromosome of CS (*46*). This work also showed the up-regulation of multiple *GLP* gene members in response to powdery mildew infection. Additionally, recent studies have demonstrated the regulation of crop traits (e.g., cotton fiber length and rice acclimation to UV-B radiation) by *GLP* genes (*47, 48*). Overall, there is increasing evidence for the multifaceted roles played by GLPs in plant growth and development as well as responses to abiotic and biotic stresses. Therefore, it is worthwhile to study the functions of the highly duplicated *GLP* genes on 4BL in wheat growth and response to environmental factors in the future.

From the whole chromosome arm comparisons depicted in Fig. 7A, it is clear that the 1RS translocation in Zhou8425B is unique from that present in other wheat cultivars (Aikang 58 and Kenong 199) or diploid rye lines (Lo7 and Weining). The 1RS in Zhou8425B is not only longer and but also possesses an extra segment (∼20.5 Mb) rich in three types of short tandem repeats (Fig. 7A, B). Although the function of this extra segment remains to be investigated, it can serve as a marker for tagging Zhou8425B 1RS during wheat breeding. Zhou8425B 1RS is also distinct in having expanded genes encoding AP2/ERF-ERF or B3 TFs, which have been reported to regulate plant growth, development, and responses to environmental stimuli. The B3 gene TraesZ8425B1RS01G036200, preferentially expressed in developing grains (Fig. 8D) with numerous co-expressed partners (Fig. 4D), offers a valuable opportunity for understanding the regulation of seed related processes and traits by an alien gene (from rye) in common wheat background. Considering that there are still worldwide interests in developing diverse 1RS translocations for wheat improvement (*36–39*), the 1RS sequence of Zhou8425B, as well as the 1RS analysis data generated by us, may benefit the progress of such research efforts.

By combining GWAS analysis and bi-parental mapping, this work mined a new APR locus (*YrZH3BS*) against YR disease, and further mapped it to a small interval (1 - 2 Mb) on chromosome 3BS with the aid of Zhou8425B genome sequence (Fig. 9). Pedigree and DNA marker analyses suggest that *YrZH3BS* locus in the YR resistant cultivar Xinhuamai 818 was derived from Zhou8425B (Fig. 9C). Consequently, Zhou8425B has now at least three known APR loci against YR, *YrZH22* and *YrZH84* identified by previous research and *YrZH3BS* discovered in this work. From analyzing YR disease severity (i.e., FLDS) and yield related traits of 212 wheat cultivars, it becomes clear that single or combined presence of *YrZH22*, *YrZH84* and *YrZH3BS* is associated with significantly decreased susceptibility to YR infection and enhanced yield performance, with triple combination (*YrZH22* + *YrZH84* + *YrZH3BS*) having the most pronounced effects (Tables 5 and 6). Curiously, *YrZH22*, *YrZH84* and *YrZH3BS* consistently enhanced GNPS, TGW, and GY but not SNPP (Tables 5 and 6). We speculate that these genes may act after wheat tillering stage to reduce YR severity. Further molecular cloning and functional studies of *YrZH22*, *YrZH84* and *YrZH3BS* will clarify this question, and the availability of Zhou8425B genome sequence will help to speed up these studies. Meanwhile, Zhou8425B and its derivative cultivars carrying *YrZH22*, *YrZH84* and *YrZH3BS* can be used as donors in the breeding of new wheat lines with durable resistance to YR disease. We noted that *YrZH22*, *YrZH84* and *YrZH3BS* have been efficiently transmitted from Zhou8425B to Zhoumai 22 and the varieties derived from Zhoumai 22 in past breeding work (Fig. S7). The efficiency of pyramiding *YrZH22*, *YrZH84* and *YrZH3BS* in wheat breeding can now be largely improved through marker-assisted gene transfer using with the DNA markers developed by previous research and our current work (Table S20).

In summary, we have constructed a highly contiguous genome assembly for the key wheat breeding germplasm Zhou8425B, which has elite agronomic traits and carries beneficial alleles of a wide array of genes and QTLs regulating plant height, photoperiod sensitivity, APR against YR and LR diseases, and grain yield related traits. Zhou8425B genome has unique chromosomal structure changes and its 1RS translocation is distinct from that in other hexaploid wheat cultivars (Aikang 58 and Kenong 9204) and diploid rye lines (Lo7 and Weining) with genome information. Use of Zhou8425B genome sequence has facilitated the mining and mapping of a new APR locus (*YrZH3BS*) against YR. Combination of *YrZH3BS* with two previously reported APR loci (*YrZH22* and *YrZH84*) is very effective in decreasing YR severity and promoting wheat grain yield. With its high quality in chromosome sequence, Zhou8425B genome assembly may find important applications in studying wheat genome biology and improving wheat performance through genomics-assisted breeding in the future.

## Acknowledgements

This work was supported by the Ministry of Science and Technology of China (2021YFF1000200), the National Science Foundation of China (32372132), the Science and Technology funds of Zhengzhou city (To Guozhang Kang, Guangwei Li, Jizhou Zheng, Dengbin Gu, and Tiancun Zheng), and China Postdoctoral Funds (to Yuxin Yang, GZC20230727). We thank Professors Fei Lu, Zhiyong Liu, Yijing Zhang, and Jizeng Jia for constructive advice and discussions.

## Author Contributions

D.W., G.L. and G.Y. configured and designed the project. G.L., Y.Y., M.C. and Z.W. sequenced and analyzed the genome. K.Z., C.L., X.L., and Z.H. performed molecular marker analysis of elite genes. S.C., Y.L., C.D. and G.Y. analyzed the effects of three APR genes on decreasing YR disease severity and promoting grain yield. T.Z., J.Z. and G.Y. developed Zhou8425B and evaluated its agronomic traits. X.Z., J.T., T.Z., J.Z., Y.P., D.G. and G.Y. curated the seeds of Zhou8425B derivative varieties. Y.R., X.S. and F.C. performed GWAS experiment and fine mapping of *YrZH3BS*. K.Z., G.L., H.Z. and C.S. investigated gluten gene loci and accumulation of gluten proteins in the seeds. D.G., T.Z., G.L., G.K., Y.P., Y.Y., X.Z. and D.W. obtained funds for the project. D.W., G.L. and Y.Y. designed the figures and tables and wrote the manuscript. All authors read and approved the manuscript.

## Methods

### Genome sequencing and assembly

High-quality DNA was extracted from the fresh leaves at the tillering stage of Zhou8425B, which was used for preparing 15 kb PacBio HiFi libraries using the SMRTbell Express Template Prep Kit 2.0 (Pacific Biosciences, CA, USA) following the manufacture’s instruction. After quality control, SMRTbell libraries were obtained and sequenced on the PacBio Sequel II/IIe platform (Pacific Biosciences, CA, USA) to obtain the polymerase reads. The PacBio SMRT-Analysis package (https://www.pacb.com) was employed for filtering the raw polymerase reads. High-quality HiFi reads, generated by SMRTLink (version v13.0) (https://www.pacb.com/support/software-downloads) with the parameters: --min-passes=3 --min-rq=0.99 (Fig. S8), were used for genome assembly.

The draft assembly was constructed using Hifiasm (version 0.19.5) with default parameters. Subsequently, the following steps were implemented to eliminate low-quality or contaminant sequences: (1) Use of purge_dup (version 1.2.5) (https://github.com/dfguan/purge_dups) to eliminate redundancy of HiFi reads with the following parameters: -2 -T changed.cutoffs -l XXX -m XXX -u XXX; (2) Alignment of sequences to the Nucleotide Sequence Database (NT) (ftp://ftp.ncbi.nlm.nih.gov/blast/db/) to filter out contaminate sequence; (3) Calculation of contig sequence depth and GC content, filtering out contigs with coverage depth below 5 or GC content above 50%. The resultant assembly was assessed in three ways. First, HiFi reads were aligned to the assembly using minimap2 (version 2.26) (https://github.com/lh3/minimap2) with the parameters: -I6G -ax map-pb -t 60, achieving a mapping rate of 99.96%. Second, based on the above alignment result, samtools (version 1.17) and bedtools (version 2.2.29) were used to calculate depth and GC content in a given window (10 kb). The GC-Depth was plotted using R (https://www.r-project.org/). Third, the assembly was evaluated using the BUSCO (version 5.1.3) tool (https://busco.ezlab.org/) with parameters: -c 60 -m geno -offline.

### Hi-C assisted scaffolding

The Hi-C libraries were constructed following established protocols (*49, 50*), which were sequenced on the Illumina NovaSeq 6000 platform. To obtain high quality Hi-C reads, the raw Hi-C sequencing data were filtered in three steps: (1) removal of reads with 3 unidentified nucleotides; (2) elimination of reads aligned to the adaptor; (3) exclusion of reads with ≥ 20% bases having phred quality ≤ 5.

The filtered Hi-C reads were mapped to the HiFi genome assembly using Bowtie2 (version 2.3.5). The 3D-DNA software (version 180419) was employed to obtain the uniquely mapped and valid paired-end reads, which were then used to produce chromosome level scaffolds with Juicebox (version 1.11.08). HiCExplorer (version 3.7.2) was utilized to generate the contact map.

### Genome annotation

Repetitive element prediction utilized a hybrid approach combining *de novo* and homology-based methods, with different types of repeats identified and annotated as described in a previous study (*28*). To annotate gene structures, a comprehensive approach involving *ab initio* prediction, protein-based homology searches, and annotation with RNA sequencing data was employed (*28*). For functional annotation, protein-coding genes were searched against three integrated protein databases: NR, SwissProt, and eggNOG. InterPro annotated protein domains and Gene Ontology terms were obtained from the corresponding eggnog-mapper annotation entry for each gene. Pathway assignments were made by BLAST against the KEGG database. The tRNAscan-SE software (version 4.09) was employed for predicting tRNAs. Other types of ncRNA were annotated using the Pfam database through BLAST (version 2.12.0) with default parameters.

### RNA sequencing

Total RNAs, extracted from 12 Zhou8425B plant samples (Table S6), were subjected to RNA sequencing as reported before (*28*). Clean reads were aligned to Z8425B genome assembly using Hisat2 (version 2.5.3a), with transcript assembly facilitated by Stringtie (version 1.3.3.b). The expression level, represented by FPKM, was calculated using the R package Ballgown. Weighted gene co-expression network analysis was executed using the R package WGCNA (https://cran.r-project.org/web/packages/WGCNA/index.html), with the Cytoscape software (version 3.10.1) used to visualize co-expression networks.

### Analysis of structure variations of Zhou8425B genome

The Python version of JCVI software (https://github.com/tanghaibao/jcvi) was used to investigate genome collinearity between Zhou8425B and 16 previously sequenced wheat varieties for revealing the unique translocations in Zhou8425B. To calculate the impact of translocation on gene expression, we extracted genes within the Zhou8425B unique translocations, and calculated the expression levels of these genes in four tissues of Zhou8425B and Aikang 58. The gene expression data of Aikang 58 had been published before (*9*). The softwares Mummer (version 3.20) and SyRI (version 1.5) were used to investigate collinearity relationships among the five 1RS chromosome arms in Zhou8425B, Aikang 58, Kenong 9204, Lo7, and Weining to uncover their structural similarities and differences.

### Investigation of transcription factor genes and gene families carried by 1RS

To explore the TF genes in 1RS and 1BS chromosome arms, we extracted the protein sequences of 1RS/1BS and used the iTAK pipeline (http://bioinfo.bti.cornell.edu/tool/itak) for annotating TFs. We further calculated the expanded TF families in Zhou8425B and examined their expression levels using transcriptome sequencing data generated with the 12 samples of Zhou8425B. The Orthofinder (version 2.3.8) program was used to perform gene family analysis for the 1RS arms in Zhou8425B, Aikang 58, Kenong 9204, Lo7, and Weining. The shared and unique gene families were computed using the online software Jvenn (https://jvenn.toulouse.inra.fr/app/example.html).

### Detection of tandemly duplicated gene clusters

The MCScanX software was used to compute TDGCs in Zhou8425B and 16 previously sequenced wheat cultivars. The online CD-search tool (https://www.ncbi.nlm.nih.gov/Structure/bwrpsb/bwrpsb.cgi) was used to discover conserved protein domains in TDGC paralogs. The three largest TDGCs in Zhou8425B were selected for further investigation. The expression levels of their paralogs were calculated using the transcriptome sequencing data generated with 12 Zhou8425B plant samples (Table S6).

### Identification and fine mapping of *YrZH3BS*

GWAS analysis was performed using the FLDS data of 245 wheat cultivars grown in five different field environments with natural and artificially induced YR infections (*14*). Genotyping of the 245 cultivars was accomplished using the 660K wheat SNP array (*51*). Bi-parental mapping of *YrZH3BS* used the F_2:3_ lines prepared with Xinhuamai 818 and Yumai 1 as parents. FLDS was scored for F_2:3_ lines and their parents as described above. Polymorphic DNA markers (Table S18) were developed for the *YrZH3BS* region using indels revealed by aligning the 3BS sequences of Zhou8425B and CS.

### Analysis of the effects of *YrZH3BS*, *YrZH22*, and *YrZH84*

A total of 212 Zhou8425B derivative varieties (Table S19) were grown in two field environments with natural and artificially induced YR infections (*14*), with their FLDS data collected as above. Four yield related traits, i.e., SNPP, GNPS, TGW, and GY, were also scored for these lines. Presence of *YrZH3BS*, *YrZH22*, and *YrZH84* in the 212 cultivars were identified using DNA markers (Table S20). The resultant genotyping information was employed to examine the effects of *YrZH3BS*, *YrZH22*, and *YrZH84* on decreasing FLDS and promoting yield related traits.

### Investigation of 17 agronomic trait genes in the wheat materials involved in the breeding of Zhou8425B

The six wheat lines used for developing Zhou8425B (Fig. 1A) were examined for the presence of four dwarf genes, one photoperiod insensitivity gene, five APR loci, and seven grain weight associated genes (Table 1) by DNA marker analysis as described before (*26, 30*). The PCR primers used were listed in Table S20.

## Data availability

The genome assembly and transcriptome datasets of Zhou8425B generated in this work have been submitted to the NCBI database, with the accession numbers being PRJNA1057534 and PRJNA1059052, respectively.

