## Supplemental Figures for "A highly contiguous hexaploid wheat genome assembly facilitates analysis of 1RS translocation and mining of a new adult plant resistance locus to yellow rust disease"

### Supplementary Figures

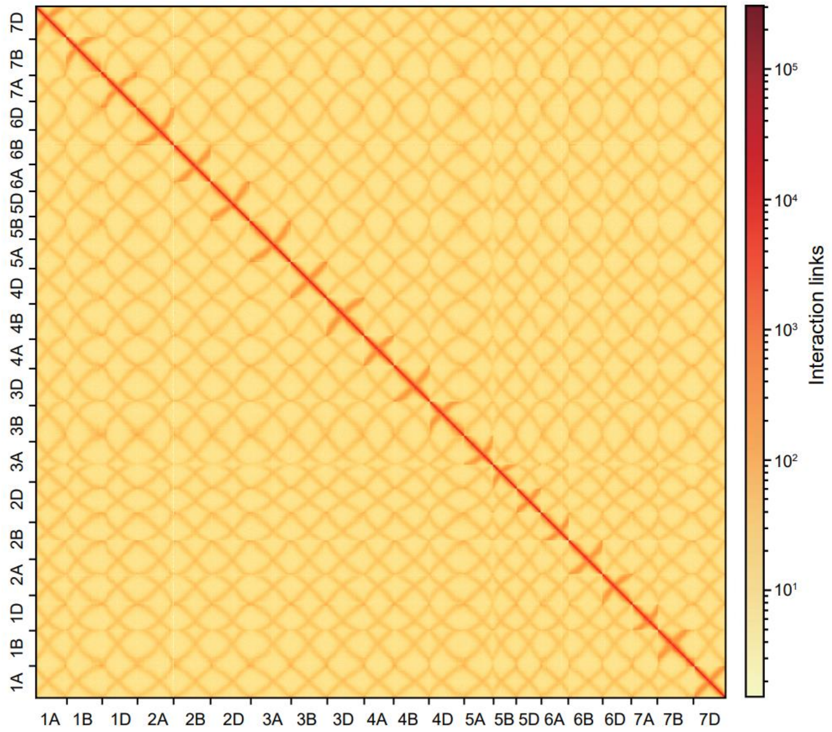


**Figure S1**. **Hi-C interaction matrix of the 21 assembled chromosomes (1A - 7 D) of Zhou8425B.** Strong intrachromosomal contacts were detected, whereas interchromosomal contacts were relatively weak. The color intensity represents frequency of contact between two 1 Mb bins.

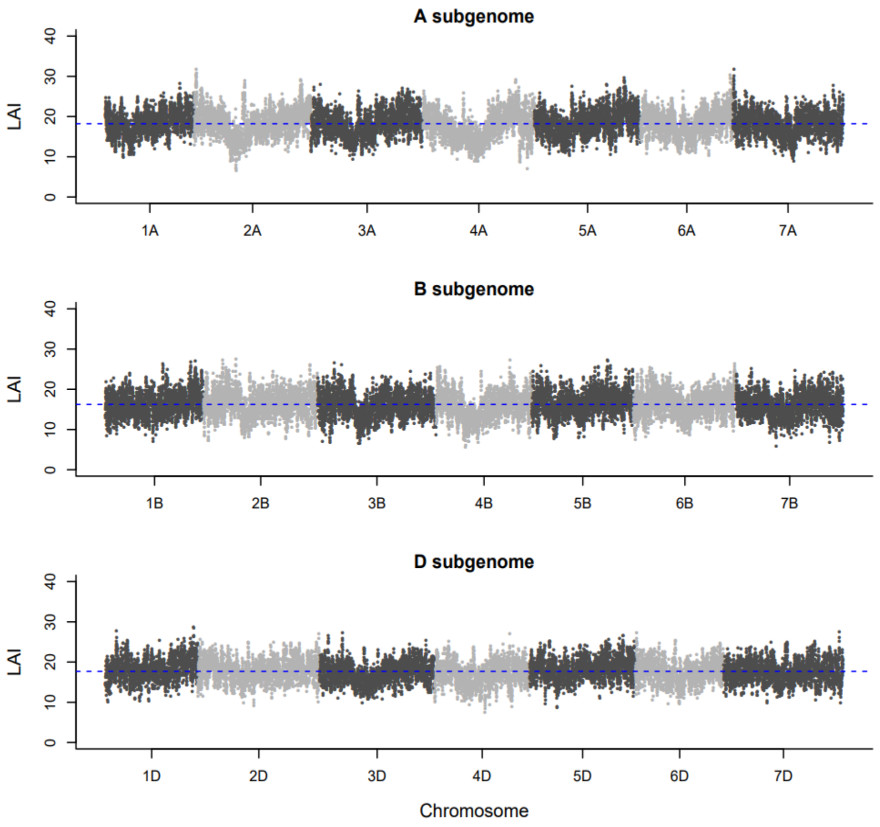


**Figure S2**. **LTR assembly index (LAI) of the three assembled subgenomes of Zhou8425B.** LAI values were computed using the method described by Ou et al. (2018). The X-axes indicate the chromosomes of A, B, and D subgenomes. Y-axes mark LAI scores of 3 Mb regions with 300 kb sliding windows on each chromosome. The average LAI values of A, B, and D subgenome (represented by the blue lines) are 18.22, 16.33, and 17.67, respectively.

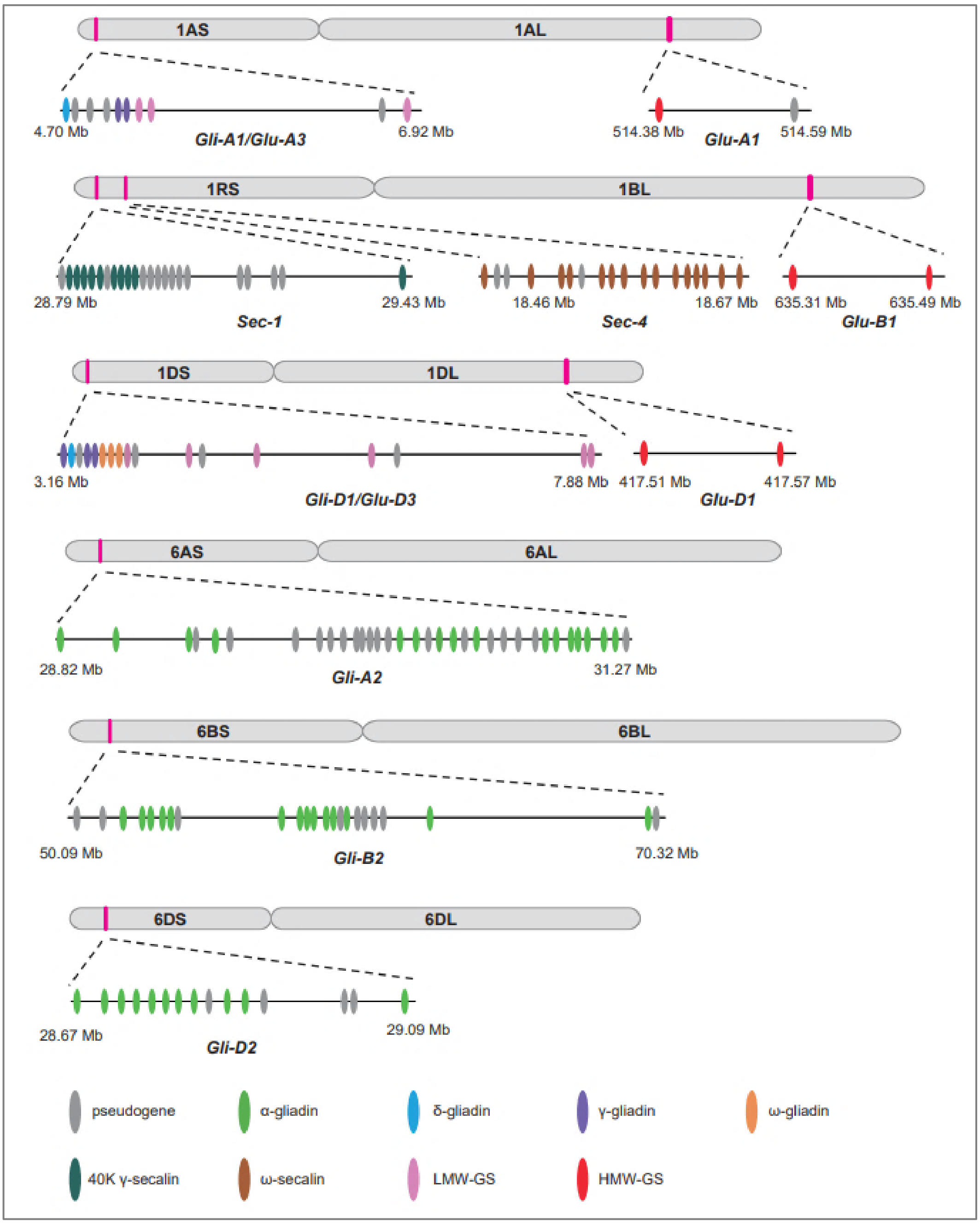


**Figure S3**. **Analysis of the gluten gene loci in Zhou8525B genome assembly.**

The chromosomal location and physical size, as well as the gluten genes contained, are shown for each locus. These loci include *Gli-A1/Glu-A3* and *Gli-D1/Glu-D3* on short arms of group 1 chromosomes, *Sec-1* and *Sec-4* on 1RS, *Glu-A1*, *-B1*, and *-D1* on long arms of group 1 chromosomes, and *Gli-A2*, *-B2*, and *-D2* on short arms of group 6 chromosomes.

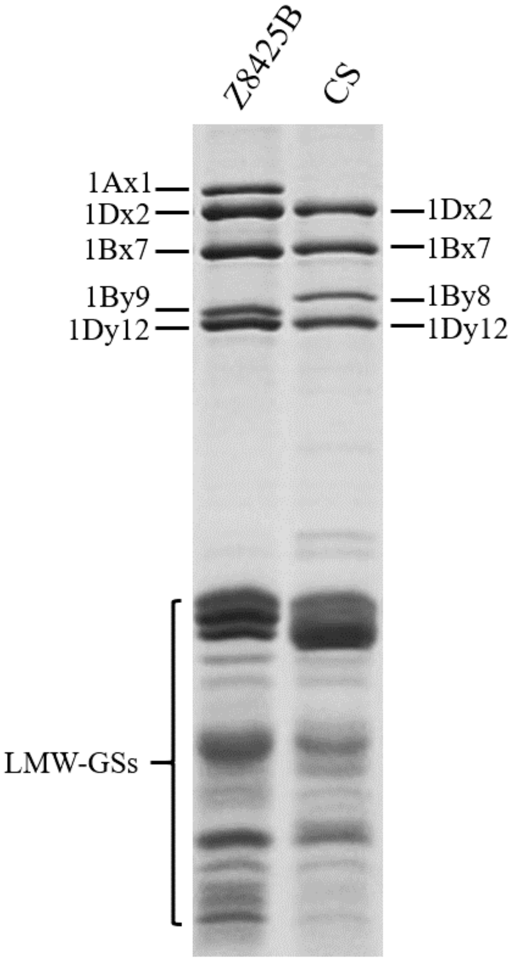


**Figure S4**. **SDS-PAGE detection of HMW-GSs and LMW-GSs in Zhou8425B mature grains.** The high-molecular-weight glutenin subunits (HMW-GSs) and low-molecular-weight glutenin subunits (LMW-GSs) extracted from Chinese Spring (CS) mature grains were used as controls. Five HMW-GSs (1Ax1, 1Bx7, 1By9, 1Dx2, and 1Dy12) and multiple LMW-GSs were accumulated in Zhou8425B (Z8425B) grains.

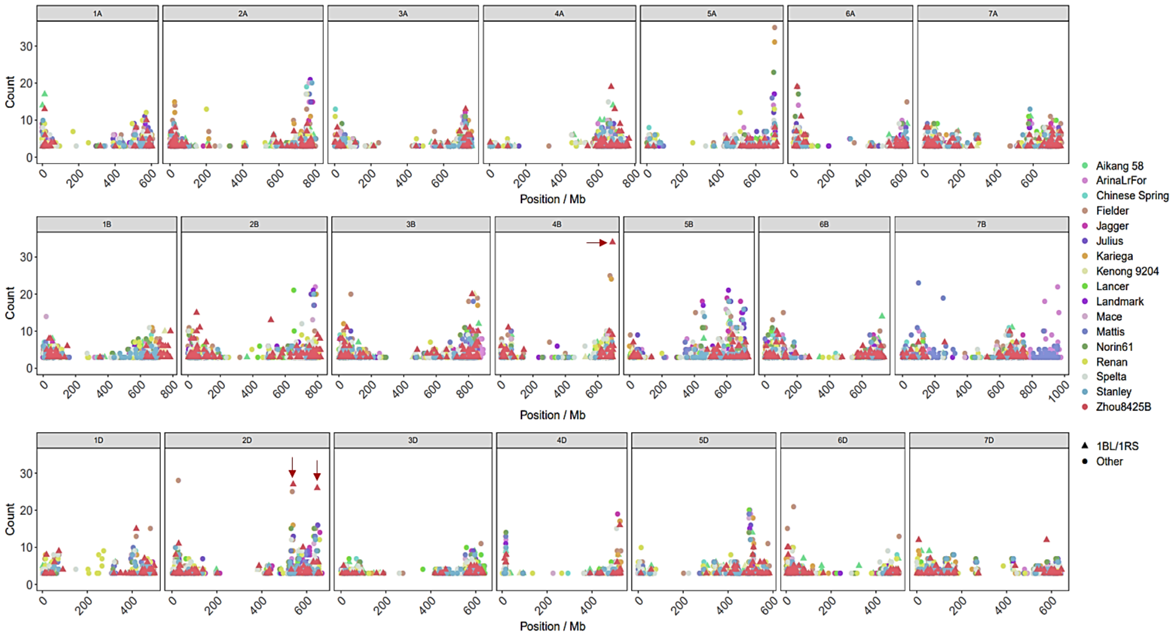


**Figure S5**. **Computation of tandemly duplicated gene clusters.**

The genome assembly of Zhou8425B and those of 16 previously sequenced common wheat varieties were analyzed using the 'duplicate_gene_classifier' program in the MCScanX package. The tandemly duplicated gene clusters (TDGCs) detected for each chromosome were shown according to their physical locations (X axes). The Y axes indicate number of paralogs in the clusters. The three super TDGCs found on the 4BL and 2DL of Zhou8425B are labeled with red arrows.

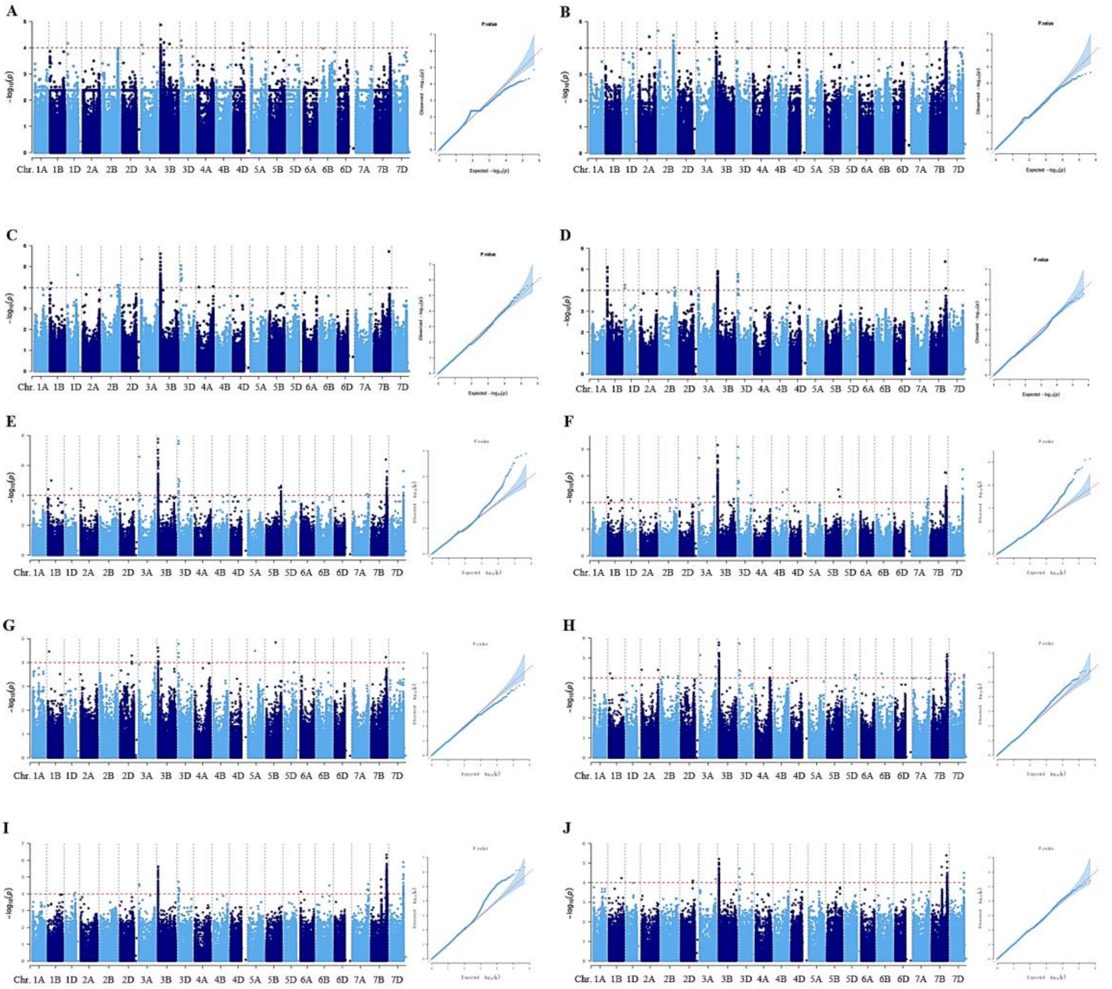


**Figure S6**. **Manhattan and QQ plots obtained by GWAS analysis with the flag leaf disease severity data of 245 common wheat varieties cultivated in five different environments with two biological repeats in each.**

**A** and **B**, two repeats at Xindu in 2017. **C** and **D**, two repeats at Xindu in 2018. **E** and **F**, two repeats at Pixian in 2017. **G** and **H**, two repeats at Pixian in 2018. **I** and **J**, two repeats at Pixian in 2019.

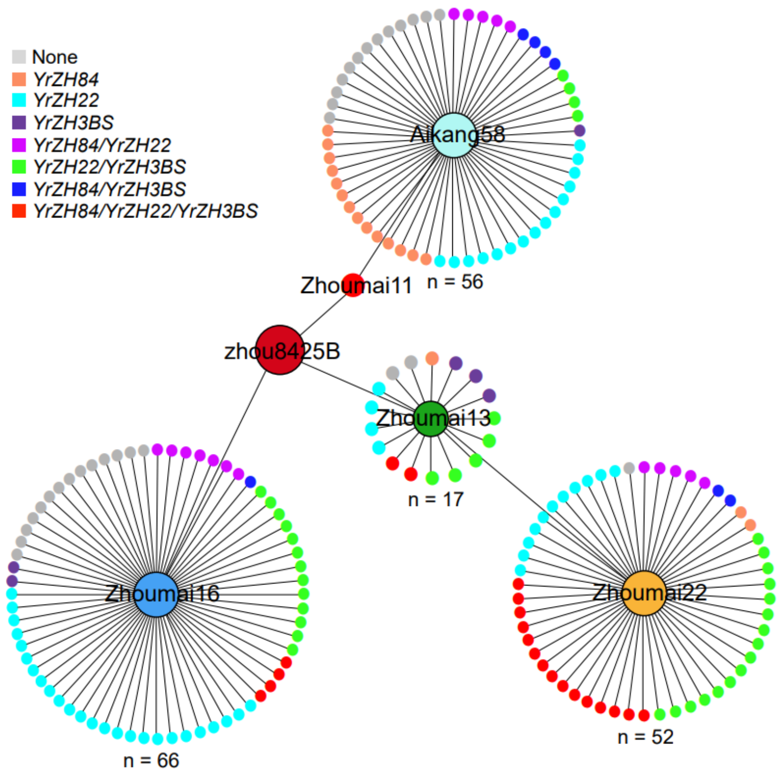


**Figure S7**. **Genetic transmission of *YrZH22*, *YrZH84* and *YrZH3BS* from Zhou8425B to its derivative cultivars.**

A subsection of Zhou8425B derivative cultivars were analyzed for the single or combined presence of three APR loci (*YrZH84*, *YrZH22*, and *YrZH3BS*) against yellow rust disease using the DNA markers listed in Table S20. Zhoumai 11, Zhoumai 13, Zhoumai 16, Zhoumai 22, and Aikang 58 are early-generation descendants of Zhou8425B. "n" indicates the number of derivative cultivars of Zhoumai 13, Zhoumai 16, Zhoumai 22, or Aikang 58. The names of the cultivars analyzed are available upon request to Dr. Shulin Chen.

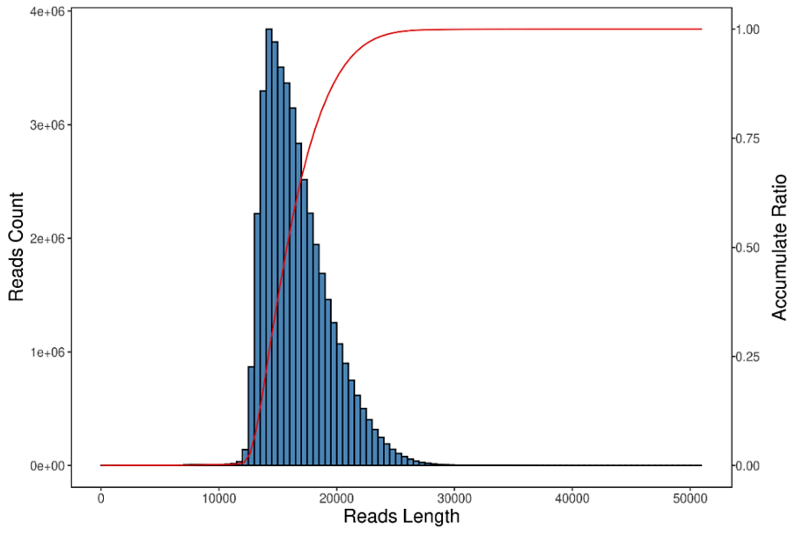


**Figure S8. Distribution of the lengths of HiFi reads generated in this study.**
The x-axis represents the length of HiFi reads. The left y-axis indicates number of HiFi reads with different lengths. The right y-axis denotes the cumulative percentage of the total bases in HiFi reads.
