## Supplemental Tables for "A highly contiguous hexaploid wheat genome assembly facilitates analysis of 1RS translocation and mining of a new adult plant resistance locus to yellow rust disease"

### Supplementary Tables

**Table S1. List of 839 Zhou8425B derivative cultivars that passed provincial and/or national certification in China**

| Number | Cultivar name | Provincial/National Certification | Year of release |
|---|---|---|---|
| 1 | Zhoumai 13 | Henan Province Identification 1999 | 1999 |
| 2 | Zhoumai 11 | National 20000007 | 2000 |
| 3 | Zhoumai 12 | National 20000008 | 2000 |
| 4 | Zhoumai 16 | National 2003029 | 2003 |
| 5 | Zhoumai 17 | National 2004008 | 2004 |
| 6 | Fanmai 5 | National 2005007 | 2005 |
| 7 | Bainong AK58 | National 2005008 | 2005 |
| 8 | Zhoumai 19 | Henan Province 2005001 | 2005 |
| 9 | Pumai 10 | Henan Province 2006002 | 2006 |
| 10 | Hemai 1 | Henan Province 2006004 | 2006 |
| 11 | Zhengnong 17 | Henan Province 2006007 | 2006 |
| 12 | Zhoumia 20 | Henan Province 2006009 | 2006 |
| 13 | Yuzhan 4 | Henan Province 2006016 | 2006 |
| 14 | Yuanyu 3 | Henan Province 2006017 | 2006 |
| 15 | 04 Zhong 36 | Henan Province 2006019 | 2006 |
| 16 | Chuanyu 21 | Sichuan Province 2007007 | 2007 |
| 17 | Zhoumai 22 | National 2007007 | 2007 |
| 18 | Zhoumai 21 | National 2007013 | 2007 |
| 19 | Xumai 30 | Jiangsu Province 200704 | 2007 |
| 20 | Dengmai 996 | Henan Province 2007008 | 2007 |
| 21 | Jinmai 8 | National 2008006 | 2008 |
| 22 | Luomai 9 | National 2008007 | 2008 |
| 23 | Zhoumai 23 | National 2008008 | 2008 |
| 24 | Luomai 22 | Henan Province 2008001 | 2008 |
| 25 | Chuanyu 24 | Sichuan Province 2009005 | 2009 |
| 26 | Luomai 21 | National 2009006 | 2009 |
| 27 | Huaimai 28 | National 2009009 | 2009 |
| 28 | Zhoumai 24 | Henan Province 2009002 | 2009 |
| 29 | Zhoumai 27 | National 2011003 | 2011 |
| 30 | Fengdecunmai 1 | National 2011004 | 2011 |
| 31 | Xinong 822 | Shaanxi Province 2011001 | 2011 |
| 32 | Xuke 316 | Henan Province 2011001 | 2011 |
| 33 | Xun 2016 | Henan Province 2011004 | 2011 |
| 34 | Zhoumai 25 | Henan Province 2011018 | 2011 |
| 35 | Taixue 7 | Henan Province 2011019 | 2011 |
| 36 | Yunong 4023 | Henan Province 2011029 | 2011 |
| 37 | Zhoumai 26 | National 2012006 | 2012 |
| 38 | Pingan 8 | National 2012007 | 2012 |
| 39 | Zhengmai 7698 | National 2012009 | 2012 |
| 40 | Zhongmai 895 | National 2012010 | 2012 |
| 41 | Luomai 18 | National 2012011 | 2012 |
| 42 | Fugao 1 | Shaanxi Province 2012008 | 2012 |

| Number | Cultivar name | Provincial/National Certification | Year of release |
|---|---|---|---|
| 43 | Xumai 32 | Jiangsu Province 201206 | 2012 |
| 44 | Xuke 718 | Henan Province 2012001 | 2012 |
| 45 | Zhengmai 583 | Henan Province 2012003 | 2012 |
| 46 | Nongda 1108 | Henan Province 2012004 | 2012 |
| 47 | Jiaomai 266 | Henan Province 2012006 | 2012 |
| 48 | Zhongxin 78 | Henan Province 2012008 | 2012 |
| 49 | Zhengmai 0856 | Henan Province 2012012 | 2012 |
| 50 | Pumai 26 | Henan Province 2012013 | 2012 |
| 51 | Yangguang 851 | Henan Province 2012017 | 2012 |
| 52 | Xumai 33 | National 2013008 | 2013 |
| 53 | Zhoumai 28 | National 2013009 | 2013 |
| 54 | Bainong 207 | National 2013010 | 2013 |
| 55 | Huaimai 35 | National 2013011 | 2013 |
| 56 | Xinmai 23 | National 2013016 | 2013 |
| 57 | Guangmingmai 2 | Shanghai City2013002 | 2013 |
| 58 | Qinnong 578 | Shaanxi Province 2013002 | 2013 |
| 59 | Qinmai 618 | Shaanxi Province 2013009 | 2013 |
| 60 | Xinong 318 | Shaanxi Province 2013013 | 2013 |
| 61 | Fengdecunmai 5 | National 2014003 | 2014 |
| 62 | Cunmai 8 | National 2014005 | 2014 |
| 63 | Jinmai 95 | Shanxi Province 2014002 | 2014 |
| 64 | Tianmai 863 | Shaanxi Province 2014010 | 2014 |
| 65 | Baomai 5 | Jiangsu Province 201403 | 2014 |
| 66 | Xumai 9158 | Jiangsu Province 201404 | 2014 |
| 67 | Huayu 198 | Henan Province 2014003 | 2014 |
| 68 | Jindi 828 | Henan Province 2014006 | 2014 |
| 69 | Zhongyu 9307 | Henan Province 2014007 | 2014 |
| 70 | Zhengyu 8 | Henan Province 2014008 | 2014 |
| 71 | Yanfeng 21 | Henan Province 2014009 | 2014 |
| 72 | Xinmai 9 | Henan Province 2014012 | 2014 |
| 73 | Zhengmai 314 | Henan Province 2014013 | 2014 |
| 74 | Kaimai 22 | Henan Province 2014017 | 2014 |
| 75 | Xuke 168 | Henan Province 2014020 | 2014 |
| 76 | Wenmai 28 | Henan Province 2014021 | 2014 |
| 77 | Zhengyumai 043 | Henan Province 2014023 | 2014 |
| 78 | Xinmai 30 | Henan Province 2014026 | 2014 |
| 79 | Xinong 188 | Shaanxi Province 2015004 | 2015 |
| 80 | Xian 240 | Shaanxi Province 2015005 | 2015 |
| 81 | Xingmin 618 | Shaanxi Province 2015008 | 2015 |
| 82 | Taimai 733 | Shaanxi Province 2015012 | 2015 |
| 83 | Xinong 805 | Shaanxi Province 2015018 | 2015 |
| 84 | Nongmai 1 | Jiangsu Province 201504 | 2015 |
| 85 | Huaimai 39 | Jiangsu Province 201507 | 2015 |
| 86 | Guomai 9 | Anhui Province 2015003 | 2015 |
| 87 | Xunda 104 | Henan Province 2015002 | 2015 |
| 88 | Zaoxiang 158 | Henan Province 2015003 | 2015 |
| 89 | Xunda 106 | Henan Province 2015008 | 2015 |
| 90 | Yanke 048 | Henan Province 2015017 | 2015 |

| Number | Cultivar name | Provincial/National Certification | Year of release |
|---|---|---|---|
| 91 | Jinsui 116 | Henan Province 2015018 | 2015 |
| 92 | Mengmai 023 | Henan Province 2015021 | 2015 |
| 93 | Lantian 35 | Gansu Province 2016014 | 2016 |
| 94 | Zhoumai 30 | National 2016006 | 2016 |
| 95 | Deyan 8 | National 2016007 | 2016 |
| 96 | Guanmai 1 | National 2016008 | 2016 |
| 97 | Luomai 29 | National 2016009 | 2016 |
| 98 | Baomai 6 | National 2016010 | 2016 |
| 99 | Xuke 129 | National 2016011 | 2016 |
| 100 | Zhengmai 379 | National 2016013 | 2016 |
| 101 | Zhengpinmai 8 | National 2016014 | 2016 |
| 102 | Shengyuan 619 | National 2016015 | 2016 |
| 103 | Zhongyu 1123 | National 2016019 | 2016 |
| 104 | Zhongyuan 18 | National 2016020 | 2016 |
| 105 | Hongdi 95 | Shandong Province 2016008 | 2016 |
| 106 | Hongdi166 | Shandong Province 20160061 | 2016 |
| 107 | Qimin 7 | Shandong Province 20160062 | 2016 |
| 108 | Mingmai 16 | Jiangsu Province 20160005 | 2016 |
| 109 | Xunong 029 | Jiangsu Province 20160007 | 2016 |
| 110 | Longke 1109 | Anhui Province 2016001 | 2016 |
| 111 | Guomai 11 | Anhui Province 2016003 | 2016 |
| 112 | Anke 0817 | Anhui Province 2016012 | 2016 |
| 113 | Lvyu 7 | Anhui Province 2016025 | 2016 |
| 114 | Huiyan 22 | Anhui Province 2016029 | 2016 |
| 115 | Huachengmai 1688 | Anhui Province 2016030 | 2016 |
| 116 | Wanmai 203 | Anhui Province 2016036 | 2016 |
| 117 | Huamai 1168 | Hubei Province 2017001 | 2017 |
| 118 | Hengjinmai 8 | National 20170002 | 2017 |
| 119 | Xumai 35 | National 20170007 | 2017 |
| 120 | Quanmai 890 | National 20170008 | 2017 |
| 121 | Puxing 5 | National 20170009 | 2017 |
| 122 | Fengdecunmai 12 | National 20170010 | 2017 |
| 123 | Wodemai 365 | National 20170011 | 2017 |
| 124 | Xinmai 29 | National 20170012 | 2017 |
| 125 | Yangao 21 | National 20170013 | 2017 |
| 126 | Mengmai 028 | Shaanxi Province 2017004 | 2017 |
| 127 | Qianmai 088 | Jiangsu Province 20170004 | 2017 |
| 128 | Jiangmai 23 | Jiangsu Province 20170005 | 2017 |
| 129 | Baomai 330 | Jiangsu Province 20170006 | 2017 |
| 130 | Liumai 618 | Anhui Province 2017002 | 2017 |
| 131 | Xihong 3 | Anhui Province 2017021 | 2017 |
| 132 | Wangjiamai 2 | Anhui Province 2017022 | 2017 |
| 133 | Yangao 58 | Henan Province 2017004 | 2017 |
| 134 | Luomai 31 | Henan Province 2017009 | 2017 |
| 135 | Changmai 9 | Henan Province 2017010 | 2017 |
| 136 | Tunmai 127 | Henan Province 2017011 | 2017 |
| 137 | Huayumai 1 | Henan Province 2017012 | 2017 |
| 138 | Tunmai 128 | Henan Province 2017016 | 2017 |

| Number | Cultivar name | Provincial/National Certification | Year of release |
|---|---|---|---|
| 139 | Bainong 201 | Henan Province 2017019 | 2017 |
| 140 | Luohan 19 | Henan Province 2017024 | 2017 |
| 141 | Baihan 207 | Henan Province 2017025 | 2017 |
| 142 | Shengmai 15 | Henan Province 2017027 | 2017 |
| 143 | Tianxuan 63 | Gansu Province 20180010 | 2018 |
| 144 | Lantian 36 | Gansu Province 20180016 | 2018 |
| 145 | Guohong 3 | National 20180006 | 2018 |
| 146 | Xinmai 32 | National 20180013 | 2018 |
| 147 | Xinnong 518 | National 20180015 | 2018 |
| 148 | Zhoumai 32 | National 20180021 | 2018 |
| 149 | Jinxiu 21 | National 20180023 | 2018 |
| 150 | Luomai 26 | National 20180025 | 2018 |
| 151 | Lunxuan 13 | National 20180026 | 2018 |
| 152 | Zhengmai 618 | National 20180027 | 2018 |
| 153 | Saidemai 1 | National 20180028 | 2018 |
| 154 | Xinkemai 169 | National 20180033 | 2018 |
| 155 | Zhongyu 1211 | National 20180035 | 2018 |
| 156 | Pumai 6311 | National 20180037 | 2018 |
| 157 | Gaomai 6 | National 20180038 | 2018 |
| 158 | Xinmai 36 | National 20180041 | 2018 |
| 159 | Zhoumai 36 | National 20180042 | 2018 |
| 160 | Zhumai 328 | National 20180047 | 2018 |
| 161 | Longmai 808 | Hebei Province 20180005 | 2018 |
| 162 | Zhongmai 3284 | Beijing City20180004 | 2018 |
| 163 | Xumai 36 | Shandong Province 20180010 | 2018 |
| 164 | Hemai 21 | Shandong Province 20180015 | 2018 |
| 165 | Qimin 9 | Shandong Province 20180021 | 2018 |
| 166 | Hongdi 176 | Shandong Province 20180026 | 2018 |
| 167 | Mengmai 032 | Shaanxi Province 2018003 | 2018 |
| 168 | Xinong 109 | Shaanxi Province 2018010 | 2018 |
| 169 | Luomai 34 | Henan Province 20180001 | 2018 |
| 170 | Xumai 318 | Henan Province 20180003 | 2018 |
| 171 | Yingman 208 | Henan Province 20180004 | 2018 |
| 172 | Tunfeng 809 | Henan Province 20180005 | 2018 |
| 173 | Zhaofeng 3188 | Henan Province 20180006 | 2018 |
| 174 | Zhengyu 11 | Henan Province 20180009 | 2018 |
| 175 | Bainong 889 | Henan Province 20180010 | 2018 |
| 176 | Zaoxiang 168 | Henan Province 20180011 | 2018 |
| 177 | Zhongzhi 0914 | Henan Province 20180012 | 2018 |
| 178 | Jinfeng 205 | Henan Province 20180013 | 2018 |
| 179 | Puxing 8 | Henan Province 20180014 | 2018 |
| 180 | Taimai 863 | Henan Province 20180015 | 2018 |
| 181 | Mengmai 0818 | Henan Province 20180017 | 2018 |
| 182 | Cunmai 18 | Henan Province 20180018 | 2018 |
| 183 | Xunmai 169 | Henan Province 20180019 | 2018 |
| 184 | Xinhuamai 818 | Henan Province 20180022 | 2018 |
| 185 | Wenyuan 0520 | Henan Province 20180026 | 2018 |
| 186 | Ping'an 0602 | Henan Province 20180027 | 2018 |

| Number | Cultivar name | Provincial/National Certification | Year of release |
|---|---|---|---|
| 187 | Fanmai 536 | Henan Province 20180028 | 2018 |
| 188 | Fengdecunmai 20 | Henan Province 20180030 | 2018 |
| 189 | Kelinmai 969 | Henan Province 20180032 | 2018 |
| 190 | Saidemai 7 | Henan Province 20180033 | 2018 |
| 191 | Yuntai 301 | Henan Province 20180034 | 2018 |
| 192 | Shunmai 6 | Henan Province 20180035 | 2018 |
| 193 | Yushengmai 21 | Henan Province 20180038 | 2018 |
| 194 | Huaichuan 101 | Henan Province 20180039 | 2018 |
| 195 | Huamai 998 | Henan Province 20180042 | 2018 |
| 196 | Zhongchuang 811 | Henan Province 20180045 | 2018 |
| 197 | Xinmai 39 | Henan Province 20180047 | 2018 |
| 198 | Cunmai 16 | National 20190007 | 2019 |
| 199 | Danmai 118 | National 20190008 | 2019 |
| 200 | Huawei 303 | National 20190010 | 2019 |
| 201 | Lianchuangmai 11 | National 20190012 | 2019 |
| 202 | Lunxuan 166 | National 20190013 | 2019 |
| 203 | Minfeng 3 | National 20190014 | 2019 |
| 204 | Ping'an 518 | National 20190016 | 2019 |
| 205 | Quanmai 29 | National 20190017 | 2019 |
| 206 | Taihemai 2 | National 20190018 | 2019 |
| 207 | Xinmai 35 | National 20190019 | 2019 |
| 208 | Xuke 918 | National 20190020 | 2019 |
| 209 | Yunong 186 | National 20190021 | 2019 |
| 210 | Zhenmai 3 | National 20190022 | 2019 |
| 211 | Zhengmai 103 | National 20190023 | 2019 |
| 212 | Zhengmai 132 | National 20190025 | 2019 |
| 213 | Zhengmai 136 | National 20190026 | 2019 |
| 214 | Zhengmai 1860 | National 20190027 | 2019 |
| 215 | Zhongnongmai 4007 | National 20190029 | 2019 |
| 216 | Zhongyu 1220 | National 20190030 | 2019 |
| 217 | Anke 1405 | National 20190032 | 2019 |
| 218 | Cunmai 11 | National 20190033 | 2019 |
| 219 | Jimai 211 | National 20190034 | 2019 |
| 220 | Saidemai 5 | National 20190035 | 2019 |
| 221 | Zhumai 305 | National 20190037 | 2019 |
| 222 | Denghai 206 | National 20190038 | 2019 |
| 223 | Zhengmai 113 | National 20190059 | 2019 |
| 224 | Yingboaifeng 8 | Hebei Province 20198016 | 2019 |
| 225 | Qimin 11 | Shandong Province 20190002 | 2019 |
| 226 | Yunhe 181 | Shandong Province 20190018 | 2019 |
| 227 | Lingmai 669 | Shaanxi Province 2019003 | 2019 |
| 228 | Xinong 388 | Shaanxi Province 2019005 | 2019 |
| 229 | Yanmai 5811 | Shaanxi Province 2019007 | 2019 |
| 230 | Xinong 836 | Shaanxi Province 2019008 | 2019 |
| 231 | Yanmai 5810 | Shaanxi Province 2019010 | 2019 |
| 232 | Xinong 9112 | Shaanxi Province 2019011 | 2019 |
| 233 | Xigao 9924 | Shaanxi Province 2019014 | 2019 |
| 234 | Datang 66 | Shaanxi Province 2019015 | 2019 |

| Number | Cultivar name | Provincial/National Certification | Year of release |
|---|---|---|---|
| 235 | Womai 102 | Anhui Province 2019001 | 2019 |
| 236 | Liumai 716 | Anhui Province 2019005 | 2019 |
| 237 | Fanyumai 20 | Henan Province 20190001 | 2019 |
| 238 | Hemai 988 | Henan Province 20190002 | 2019 |
| 239 | Nongmai 22 | Henan Province 20190004 | 2019 |
| 240 | Tunmai 259 | Henan Province 20190007 | 2019 |
| 241 | Qiule 168 | Henan Province 20190008 | 2019 |
| 242 | Zhengmai 1354 | Henan Province 20190009 | 2019 |
| 243 | Zhongyu 1526 | Henan Province 20190010 | 2019 |
| 244 | Chenbo 998 | Henan Province 20190011 | 2019 |
| 245 | Fanyumai 18 | Henan Province 20190012 | 2019 |
| 246 | Tianmin 304 | Henan Province 20190015 | 2019 |
| 247 | Baimai 1811 | Henan Province 20190016 | 2019 |
| 248 | Keda 668 | Henan Province 20190017 | 2019 |
| 249 | Hongzhan 628 | Henan Province 20190018 | 2019 |
| 250 | Jiyanmai 10 | Henan Province 20190021 | 2019 |
| 251 | Chuangxing 26 | Henan Province 20190022 | 2019 |
| 252 | Zhengpinmai 26 | Henan Province 20190023 | 2019 |
| 253 | Wenmai 968 | Henan Province 20190024 | 2019 |
| 254 | Zhuomai 6 | Henan Province 20190026 | 2019 |
| 255 | Jinzhan 638 | Henan Province 20190027 | 2019 |
| 256 | Xuyanmai 3 | Henan Province 20190028 | 2019 |
| 257 | Qifeng 5 | Henan Province 20190029 | 2019 |
| 258 | Tianmai 119 | Henan Province 20190030 | 2019 |
| 259 | Caimai 116 | Henan Province 20190031 | 2019 |
| 260 | Saidemai 6 | Henan Province 20190033 | 2019 |
| 261 | Zhengke 137 | Henan Province 20190035 | 2019 |
| 262 | Yunong 516 | Henan Province 20190036 | 2019 |
| 263 | Gaomai 8 | Henan Province 20190037 | 2019 |
| 264 | Xuyanmai 4 | Henan Province 20190038 | 2019 |
| 265 | Huamai 999 | Henan Province 20190041 | 2019 |
| 266 | Luofeng 168 | Henan Province 20190042 | 2019 |
| 267 | Huayu 166 | Henan Province 20190044 | 2019 |
| 268 | Xindifeng 168 | Henan Province 20190046 | 2019 |
| 269 | Caiyuan 2 | Henan Province 20190048 | 2019 |
| 270 | Xianmai 19 | Henan Province 20190049 | 2019 |
| 271 | Hemai 1310 | Henan Province 20190052 | 2019 |
| 272 | Wanmai 66 | Henan Province 20190054 | 2019 |
| 273 | Luomai 26 | National 20200008 | 2020 |
| 274 | Guanmai 2 | National 20200010 | 2020 |
| 275 | Luomai 27 | National 20200013 | 2020 |
| 276 | Aimai 24 | National 20200017 | 2020 |
| 277 | Xumai 511 | National 20200018 | 2020 |
| 278 | Taihemai 5 | National 20200023 | 2020 |
| 279 | Zhengpinmai 25 | National 20200024 | 2020 |
| 280 | Huawei 307 | National 20200025 | 2020 |
| 281 | Qimin 8 | National 20200032 | 2020 |
| 282 | Pumai 168 | National 20200044 | 2020 |

| Number | Cultivar name | Provincial/National Certification | Year of release |
|---|---|---|---|
| 283 | Zhengmai 6694 | National 20200045 | 2020 |
| 284 | Xumai 178 | National 20200047 | 2020 |
| 285 | Zhongmai 875 | National 20200056 | 2020 |
| 286 | Yongfeng 101 | National 20200059 | 2020 |
| 287 | Womai 1211 | National 20200061 | 2020 |
| 288 | Zhongyu 9302 | National 20200062 | 2020 |
| 289 | Jiangmai 186 | National 20200065 | 2020 |
| 290 | Huiyan 912 | National 20200067 | 2020 |
| 291 | Shangmai 156 | National 20200069 | 2020 |
| 292 | Xumai 2023 | National 20200071 | 2020 |
| 293 | Fanmai 803 | National 20200073 | 2020 |
| 294 | Xinkemai 168 | National 20200075 | 2020 |
| 295 | Xinong 100 | National 20200076 | 2020 |
| 296 | Xinong 99 | National 20200077 | 2020 |
| 297 | Jixing 653 | National 20200078 | 2020 |
| 298 | Yuanfeng 369 | National 20200079 | 2020 |
| 299 | Jinli 9 | National 20200082 | 2020 |
| 300 | Qiule 6 | National 20200085 | 2020 |
| 301 | Jinchengmai 17 | National 20200086 | 2020 |
| 302 | Pingmai 189 | National 20200090 | 2020 |
| 303 | Pumai 053 | National 20200091 | 2020 |
| 304 | Zhoumai 33 | National 20200092 | 2020 |
| 305 | Luomai 28 | National 20200093 | 2020 |
| 306 | Liangyuanmai 2 | National 20200095 | 2020 |
| 307 | Fengdecunmai 13 | National 20200097 | 2020 |
| 308 | Jike 667 | Hebei Province 20208014 | 2020 |
| 309 | Qimin 13 | Shandong Province 20200016 | 2020 |
| 310 | Xinxing 617 | Shandong Province 20206028 | 2020 |
| 311 | Ruihuamai 506 | Jiangsu Province 20190010 | 2020 |
| 312 | Zhongyuanfeng 1 | Henan Province 20200001 | 2020 |
| 313 | Neile 269 | Henan Province 20200003 | 2020 |
| 314 | Zhongyanmai 6 | Henan Province 20200007 | 2020 |
| 315 | Heda 518 | Henan Province 20200009 | 2020 |
| 316 | Wenhe 902 | Henan Province 20200010 | 2020 |
| 317 | Bainong 219 | Henan Province 20200013 | 2020 |
| 318 | Caizhi 566 | Henan Province 20200014 | 2020 |
| 319 | Juchengmai 6 | Henan Province 20200016 | 2020 |
| 320 | Bainong 307 | Henan Province 20200017 | 2020 |
| 321 | Wenmai 168 | Henan Province 20200018 | 2020 |
| 322 | Jindi 8931 | Henan Province 20200019 | 2020 |
| 323 | Changmai 15 | Henan Province 20200021 | 2020 |
| 324 | Caizhi 16 | Henan Province 20200022 | 2020 |
| 325 | Huake 016 | Henan Province 20200023 | 2020 |
| 326 | Xianmai 18 | Henan Province 20200025 | 2020 |
| 327 | Senke 093 | Henan Province 20200028 | 2020 |
| 328 | Fumai 916 | Henan Province 20200032 | 2020 |
| 329 | Zhengmai 6687 | Henan Province 20200033 | 2020 |
| 330 | Shangmai 8 | Henan Province 20200036 | 2020 |

| Number | Cultivar name | Provincial/National Certification | Year of release |
|---|---|---|---|
| 331 | Chunxiao 158 | Henan Province 20200038 | 2020 |
| 332 | Tianmai 166 | Henan Province 20200039 | 2020 |
| 333 | Xinmai 51 | Henan Province 20200040 | 2020 |
| 334 | Hemai 32 | Henan Province 20200041 | 2020 |
| 335 | Guangtai 336 | Henan Province 20200042 | 2020 |
| 336 | Hemai 11 | Henan Province 20200043 | 2020 |
| 337 | Ruixingmai 618 | Henan Province 20200044 | 2020 |
| 338 | Fumai 709 | Henan Province 20200047 | 2020 |
| 339 | Yunong 605 | Henan Province 20200049 | 2020 |
| 340 | Bainong 365 | Henan Province 20200050 | 2020 |
| 341 | Baojingmai 161 | Henan Province 20200051 | 2020 |
| 342 | Zhengjke 168 | Henan Province 20200052 | 2020 |
| 343 | Senke 267 | Henan Province 20200054 | 2020 |
| 344 | Bainong 5822 | Henan Province 20200058 | 2020 |
| 345 | Luomai 40 | Henan Province 20200061 | 2020 |
| 346 | Zhengpinmai 27 | Henan Province 20200062 | 2020 |
| 347 | Fumai 1801 | Hubei Province 20210009 | 2020 |
| 348 | Taimai 601 | Jiangsu Province 20200009 | 2020 |
| 349 | Xinong 977 | Jiangsu Province 20200014 | 2020 |
| 350 | Ruihuamai 549 | Jiangsu Province 20200018 | 2020 |
| 351 | Jiangmai 869 | Jiangsu Province 20200019 | 2020 |
| 352 | Xumai 39 | Jiangsu Province 20200030 | 2020 |
| 353 | Zhongxing 999 | Anhui Province 20210002 | 2020 |
| 354 | Guoyu 16 | Anhui Province 20210003 | 2020 |
| 355 | Jinmai 106 | Anhui Province 20210006 | 2020 |
| 356 | Guomai 203 | Anhui Province 20210007 | 2020 |
| 357 | Wanmai 1648 | Anhui Province 20210016 | 2020 |
| 358 | Guomai 103 | Anhui Province 20210018 | 2020 |
| 359 | Suyu 07021 | Anhui Province 20210035 | 2020 |
| 360 | Huacheng 3077 | Anhui Province 20210036 | 2020 |
| 361 | Yutian 2018 | Anhui Province 20211002 | 2020 |
| 362 | Fengxingmai 2 | Anhui Province 20211014 | 2020 |
| 363 | Xuyan 5 | National 20210015 | 2020 |
| 364 | Zhengmai 16 | National 20210016 | 2020 |
| 365 | Pingan 658 | National 20210020 | 2020 |
| 366 | Fengyunmai 5 | National 20210022 | 2020 |
| 367 | Baoliang 5 | National 20210028 | 2020 |
| 368 | Houdemai 981 | National 20210029 | 2020 |
| 369 | Saidemai 8 | National 20210031 | 2020 |
| 370 | Zhengmai 22 | National 20210033 | 2020 |
| 371 | Zhongyu 1428 | National 20210034 | 2020 |
| 372 | Zhumai 762 | National 20210035 | 2020 |
| 373 | Zhongmai 1818 | National 20210037 | 2020 |
| 374 | Xinong 235 | National 20210040 | 2020 |
| 375 | Xinmai 38 | National 20210042 | 2020 |
| 376 | Dehongfumai 11 | National 20210043 | 2020 |
| 377 | Chuangmai 58 | National 20210045 | 2020 |
| 378 | Huiyan 56 | National 20210064 | 2020 |

| Number | Cultivar name | Provincial/National Certification | Year of release |
|---|---|---|---|
| 379 | Guomai 505 | National 20210079 | 2020 |
| 380 | Pumai 8062 | National 20210080 | 2020 |
| 381 | Pumai 117 | National 20210081 | 2020 |
| 382 | Tianmai 160 | National 20210084 | 2020 |
| 383 | Changmai 20 | National 20210086 | 2020 |
| 384 | Zhongmai 6032 | National 20210094 | 2020 |
| 385 | Zhengmai 150 | National 20210114 | 2020 |
| 386 | Xinliang 9 | National 20210115 | 2020 |
| 387 | Zhengmai 9188 | National 20210117 | 2020 |
| 388 | Liangyuan 666 | National 20210119 | 2020 |
| 389 | Suyanmai 658 | National 20210120 | 2020 |
| 390 | Bainong 418 | National 20210121 | 2020 |
| 391 | Huaimai 178 | National 20210127 | 2020 |
| 392 | Chuangmai 68 | National 20210130 | 2020 |
| 393 | Xinzhi 9 | National 20210131 | 2020 |
| 394 | Hongmai 360 | National 20210133 | 2020 |
| 395 | Ronghua 116 | National 20210134 | 2020 |
| 396 | Anmai 1350 | National 20210136 | 2020 |
| 397 | Hangyu 19 | National 20210141 | 2020 |
| 398 | Sui 1309 | National 20210143 | 2020 |
| 399 | Luomai 37 | National 20210144 | 2020 |
| 400 | Aimai 180 | National 20210146 | 2020 |
| 401 | Qimin 12 | National 20210149 | 2020 |
| 402 | Ruihuamai 568 | National 20210047 | 2020 |
| 403 | Bainong 4199 | National 20210049 | 2020 |
| 404 | Fengdecunmai 23 | National 20210050 | 2020 |
| 405 | Fengdecunmai 21 | National 20210051 | 2020 |
| 406 | Zhongliang 38 | Gansu Province 20200007 | 2020 |
| 407 | Lantian 42 | Gansu Province 20200009 | 2020 |
| 408 | Xinong 106 | Shaanxi Province 2020001 | 2020 |
| 409 | Xinong 857 | Shaanxi Province 2020006 | 2020 |
| 410 | Xinong 619 | Shaanxi Province 2020007 | 2020 |
| 411 | Xianmai 519 | Shaanxi Province 2020010 | 2020 |
| 412 | Datang 63 | Shaanxi Province 2020012 | 2020 |
| 413 | Shandao 198 | Shaanxi Province 2020013 | 2020 |
| 414 | Xinong 286 | Shaanxi Province 2020023 | 2020 |
| 415 | Denghai 208 | Anhui Province 20200001 | 2020 |
| 416 | Chengmai 1599 | Anhui Province 20200009 | 2020 |
| 417 | Jinhe 991 | Hebei Province 20210021 | 2021 |
| 418 | Jinmai 109 | Shanxi Province 20200001 | 2021 |
| 419 | Jimai 108 | Shandong Province 20210009 | 2021 |
| 420 | Shannong 44 | Shandong Province 20210016 | 2021 |
| 421 | Qingzhao 19 | Shandong Province 20216036 | 2021 |
| 422 | Haomai 18 | Shandong Province 20216041 | 2021 |
| 423 | Caimai 08 | Shandong Province 20216049 | 2021 |
| 424 | Xinong 926 | Shaanxi Province 20210002 | 2021 |
| 425 | Xinong 33 | Shaanxi Province 20210004 | 2021 |
| 426 | Huamai 027 | Shaanxi Province 20210005 | 2021 |

| Number | Cultivar name | Provincial/National Certification | Year of release |
|---|---|---|---|
| 427 | Juliang 19 | Shaanxi Province 20210009 | 2021 |
| 428 | Xinong 921 | Shaanxi Province 20210011 | 2021 |
| 429 | Xinong 579 | Shaanxi Province 20210012 | 2021 |
| 430 | Xinong 116 | Shaanxi Province 20210015 | 2021 |
| 431 | Ronghua 661 | Shaanxi Province 20210018 | 2021 |
| 432 | Huaimai 66 | Jiangsu Province 20210024 | 2021 |
| 433 | Ruihuamai 563 | Jiangsu Province 20210025 | 2021 |
| 434 | Wanxinmai 799 | Anhui Province 20211015 | 2021 |
| 435 | Wanmai 818 | Anhui Province 20211018 | 2021 |
| 436 | Wanmai 303 | Anhui Province 20211020 | 2021 |
| 437 | Liumai 521 | Anhui Province 20211023 | 2021 |
| 438 | Hengmai 1618 | Anhui Province 20211024 | 2021 |
| 439 | Mengmai 2 | Anhui Province 20211025 | 2021 |
| 440 | Fengxingmai 4 | Anhui Province 20211027 | 2021 |
| 441 | Fumai 2000 | Anhui Province 20211030 | 2021 |
| 442 | Liuziheimai 1 | Anhui Province 20212003 | 2021 |
| 443 | Zhengmai 163 | Henan Province 20210001 | 2021 |
| 444 | Yanfeng 28 | Henan Province 20210002 | 2021 |
| 445 | Fumai 708 | Henan Province 20210003 | 2021 |
| 446 | Wenmai 186 | Henan Province 20210004 | 2021 |
| 447 | Zhouyumai 6 | Henan Province 20210005 | 2021 |
| 448 | Zhongyu 1702 | Henan Province 20210006 | 2021 |
| 449 | Nongmai 51 | Henan Province 20210007 | 2021 |
| 450 | Junmai 518 | Henan Province 20210008 | 2021 |
| 451 | Yanfeng 29 | Henan Province 20210009 | 2021 |
| 452 | Yanbo 369 | Henan Province 20210012 | 2021 |
| 453 | Taihe 896 | Henan Province 20210013 | 2021 |
| 454 | Zhongguomai 10 | Henan Province 20210015 | 2021 |
| 455 | Yingmao 1 | Henan Province 20210017 | 2021 |
| 456 | Wenyu 3 | Henan Province 20210018 | 2021 |
| 457 | Chenbo 3518 | Henan Province 20210020 | 2021 |
| 458 | Kemai 1609 | Henan Province 20210021 | 2021 |
| 459 | Yanke 068 | Henan Province 20210022 | 2021 |
| 460 | Zhumai 256 | Henan Province 20210023 | 2021 |
| 461 | Shuomai 895 | Henan Province 20210025 | 2021 |
| 462 | Fenghuang 619 | Henan Province 20210026 | 2021 |
| 463 | Zhengnong 4108 | Henan Province 20210030 | 2021 |
| 464 | Wanmai 788 | Henan Province 20210032 | 2021 |
| 465 | Lvyuanmai 8 | Henan Province 20210033 | 2021 |
| 466 | Luohan 30 | Henan Province 20210034 | 2021 |
| 467 | Nonghan 101 | Henan Province 20210035 | 2021 |
| 468 | Wenmai 30 | Henan Province 20210036 | 2021 |
| 469 | Yumai 198 | Henan Province 20210037 | 2021 |
| 470 | Yunong 908 | Henan Province 20210040 | 2021 |
| 471 | Zhengmai 816 | Henan Province 20210041 | 2021 |
| 472 | Yunong 902 | Henan Province 20210047 | 2021 |
| 473 | Zhengmai 179 | Henan Province 20210048 | 2021 |
| 474 | Yongheimai 1 | Henan Province 20210049 | 2021 |

| Number | Cultivar name | Provincial/National Certification | Year of release |
|---|---|---|---|
| 475 | Zhengmai 817 | Henan Province 20210060 | 2021 |
| 476 | Senke 266 | Henan Province 20210062 | 2021 |
| 477 | Chunxiao 186 | Henan Province 20210063 | 2021 |
| 478 | Tianmai 178 | Henan Province 20210065 | 2021 |
| 479 | Zhaofeng 18 | Henan Province 20210066 | 2021 |
| 480 | Xumai 1708 | Henan Province 20210069 | 2021 |
| 481 | Anmai 13 | Henan Province 20210070 | 2021 |
| 482 | Anmai 22 | Henan Province 20210071 | 2021 |
| 483 | Changmai 21 | Henan Province 20210072 | 2021 |
| 484 | Wan 1390 | Henan Province 20210073 | 2021 |
| 485 | Zhengmai 9189 | Henan Province 20210075 | 2021 |
| 486 | Zhoumai 40 | Henan Province 20210076 | 2021 |
| 487 | Cunmai 608 | Henan Province 20210077 | 2021 |
| 488 | Fannong 11 | Henan Province 20210078 | 2021 |
| 489 | Laimai 201 | Henan Province 20210080 | 2021 |
| 490 | Shenzhoumai 216 | Henan Province 20210082 | 2021 |
| 491 | Jinlishenhua 608 | Henan Province 20210084 | 2021 |
| 492 | Hemai 35 | Henan Province 20210085 | 2021 |
| 493 | Zaoxiang 208 | Henan Province 20210086 | 2021 |
| 494 | Yangguang 728 | Henan Province 20210087 | 2021 |
| 495 | Zhuomai 20 | Henan Province 20210089 | 2021 |
| 496 | Huaguan 1 | Henan Province 20210090 | 2021 |
| 497 | Huachangmai 26 | Henan Province 20210091 | 2021 |
| 498 | Yingmai 1 | Henan Province 20210092 | 2021 |
| 499 | Keda 111 | Henan Province 20210093 | 2021 |
| 500 | Pingmai 20 | Henan Province 20210094 | 2021 |
| 501 | Yunong 806 | Henan Province 20210095 | 2021 |
| 502 | Jinli 11 | Henan Province 20210097 | 2021 |
| 503 | Chuangxingmai 102 | Henan Province 20210105 | 2021 |
| 504 | Pumai 1128 | Henan Province 20210108 | 2021 |
| 505 | Wenmai 169 | Henan Province 20210109 | 2021 |
| 506 | Yujinmai 017 | Henan Province 20210110 | 2021 |
| 507 | Zhoumai 35 | Henan Province 20210112 | 2021 |
| 508 | Xinyu 178 | Henan Province 20210114 | 2021 |
| 509 | Ruixingmai 625 | Henan Province 20210116 | 2021 |
| 510 | Bainong 8822 | National 20220013 | 2022 |
| 511 | Cunmai 633 | National 20220014 | 2022 |
| 512 | Fengdecunmai 22 | National 20220015 | 2022 |
| 513 | Hefeng 3 | National 20220017 | 2022 |
| 514 | Lianfeng 707 | National 20220020 | 2022 |
| 515 | Pumai 116 | National 20220021 | 2022 |
| 516 | Ruihuamai 502 | National 20220023 | 2022 |
| 517 | Shunmai 10 | National 20220025 | 2022 |
| 518 | Shunmai 11 | National 20220026 | 2022 |
| 519 | Wenmai 801 | National 20220028 | 2022 |
| 520 | Guomai 33 | National 20220029 | 2022 |
| 521 | Xumai 44 | National 20220032 | 2022 |
| 522 | Xuke 6 | National 20220034 | 2022 |

| Number | Cultivar name | Provincial/National Certification | Year of release |
|---|---|---|---|
| 523 | Yanfeng 168 | National 20220035 | 2022 |
| 524 | Yongfeng 103 | National 20220036 | 2022 |
| 525 | Yongminmai 1 | National 20220037 | 2022 |
| 526 | Zhengmai 18 | National 20220038 | 2022 |
| 527 | Zhongzhimai 13 | National 20220039 | 2022 |
| 528 | Zhoumai 37 | National 20220040 | 2022 |
| 529 | Zhoumai 38 | National 20220041 | 2022 |
| 530 | Luohan 28 | National 20220055 | 2022 |
| 531 | Anke 1803 | National 20220072 | 2022 |
| 532 | Hengjinmai 9 | National 20220073 | 2022 |
| 533 | Wankenmai 1702 | National 20220079 | 2022 |
| 534 | Xinmai 52 | National 20220080 | 2022 |
| 535 | Xumai 41 | National 20220081 | 2022 |
| 536 | Zhengmai 1835 | National 20220082 | 2022 |
| 537 | Jimai 117 | National 20220086 | 2022 |
| 538 | Malan 11 | National 20220090 | 2022 |
| 539 | Bainong 1316 | National 20220104 | 2022 |
| 540 | Baojingmai 188 | National 20220105 | 2022 |
| 541 | Delichangmai 1 | National 20220106 | 2022 |
| 542 | Dingyan 189 | National 20220107 | 2022 |
| 543 | Guohemai 3 | National 20220110 | 2022 |
| 544 | Guoxinmai 188 | National 20220111 | 2022 |
| 545 | Huakenmai 7 | National 20220113 | 2022 |
| 546 | Huazhan 177 | National 20220114 | 2022 |
| 547 | Huamai 169 | National 20220115 | 2022 |
| 548 | Huaimai 59 | National 20220116 | 2022 |
| 549 | Jinchengmai 19 | National 20220118 | 2022 |
| 550 | Jinyongfeng 99 | National 20220119 | 2022 |
| 551 | Jinmai 35 | National 20220120 | 2022 |
| 552 | Kaimai 1701 | National 20220122 | 2022 |
| 553 | Kelin 618 | National 20220123 | 2022 |
| 554 | Luomai 36 | National 20220126 | 2022 |
| 555 | Nongmai 176 | National 20220127 | 2022 |
| 556 | Ruimai 16 | National 20220129 | 2022 |
| 557 | Tianzhongmai 7 | National 20220132 | 2022 |
| 558 | Wofengmai 168 | National 20220134 | 2022 |
| 559 | Wofengmai 169 | National 20220135 | 2022 |
| 560 | Wofengmai 188 | National 20220136 | 2022 |
| 561 | Xinong 598 | National 20220137 | 2022 |
| 562 | Xianmai 22 | National 20220140 | 2022 |
| 563 | Xinmai 40 | National 20220141 | 2022 |
| 564 | Xumai 40 | National 20220147 | 2022 |
| 565 | Yunong 916 | National 20220149 | 2022 |
| 566 | Yuanfeng 519 | National 20220150 | 2022 |
| 567 | Zhaofeng 3188 | National 20220152 | 2022 |
| 568 | Zhaofeng 6 | National 20220153 | 2022 |
| 569 | Zhengke 11 | National 20220154 | 2022 |
| 570 | Zhongmai 34 | National 20220157 | 2022 |

| Number | Cultivar name | Provincial/National Certification | Year of release |
|---|---|---|---|
| 571 | Zhongmai 5215 | National 20220158 | 2022 |
| 572 | Zhongyanmai 688 | National 20220159 | 2022 |
| 573 | Zhongzhongmai 21 | National 20220161 | 2022 |
| 574 | Liangshan 108 | National 20220170 | 2022 |
| 575 | Gufeng 8 | Henan Province 20220001 | 2022 |
| 576 | Ruiyanmai 168 | Henan Province 20220002 | 2022 |
| 577 | Wenfeng 218 | Henan Province 20220005 | 2022 |
| 578 | Zhongnong 867 | Henan Province 20220006 | 2022 |
| 579 | Yangao 161 | Henan Province 20220007 | 2022 |
| 580 | Shangmai 188 | Henan Province 20220008 | 2022 |
| 581 | Fanyumai 29 | Henan Province 20220010 | 2022 |
| 582 | Yanbo 307 | Henan Province 20220011 | 2022 |
| 583 | Baimai 5811 | Henan Province 20220012 | 2022 |
| 584 | Wenmai 2216 | Henan Province 20220013 | 2022 |
| 585 | Puxing 16 | Henan Province 20220014 | 2022 |
| 586 | Mengmai 169 | Henan Province 20220015 | 2022 |
| 587 | Zhongdimai 607 | Henan Province 20220017 | 2022 |
| 588 | Taixue 278 | Henan Province 20220018 | 2022 |
| 589 | Wennong 301 | Henan Province 20220019 | 2022 |
| 590 | Wenyu 709 | Henan Province 20220020 | 2022 |
| 591 | Xuke 3 | Henan Province 20220021 | 2022 |
| 592 | Xinzhi 610 | Henan Province 20220022 | 2022 |
| 593 | Chunxiao 159 | Henan Province 20220023 | 2022 |
| 594 | Guangtai 266 | Henan Province 20220025 | 2022 |
| 595 | Shengke 188 | Henan Province 20220026 | 2022 |
| 596 | Dengmai 298 | Henan Province 20220027 | 2022 |
| 597 | Chunfeng 51 | Henan Province 20220029 | 2022 |
| 598 | Zhengmai 175 | Henan Province 20220030 | 2022 |
| 599 | Luomai 76 | Henan Province 20220031 | 2022 |
| 600 | Fenghuang 98 | Henan Province 20220034 | 2022 |
| 601 | Xianmai 23 | Henan Province 20220036 | 2022 |
| 602 | Yanbo 1896 | Henan Province 20220037 | 2022 |
| 603 | Yufengshuo 001 | Henan Province 20220038 | 2022 |
| 604 | Yangao 159 | Henan Province 20220039 | 2022 |
| 605 | Zhengmai 918 | Henan Province 20220040 | 2022 |
| 606 | Baimai 589 | Henan Province 20220041 | 2022 |
| 607 | Yunong 903 | Henan Province 20220046 | 2022 |
| 608 | Yunong 904 | Henan Province 20220047 | 2022 |
| 609 | Yumai 8 | Henan Province 20220049 | 2022 |
| 610 | Wanmai 362 | Henan Province 20220050 | 2022 |
| 611 | Yunong 906 | Henan Province 20220052 | 2022 |
| 612 | Yunong 910 | Henan Province 20220053 | 2022 |
| 613 | Zhengmai 821 | Henan Province 20220056 | 2022 |
| 614 | Zhengmai 824 | Henan Province 20220057 | 2022 |
| 615 | Tianmai 139 | Henan Province 20220061 | 2022 |
| 616 | Wenmai 34 | Henan Province 20220063 | 2022 |
| 617 | Zhoumai 45 | Henan Province 20220065 | 2022 |
| 618 | Heping 9 | Henan Province 20220067 | 2022 |

| Number | Cultivar name | Provincial/National Certification | Year of release |
|---|---|---|---|
| 619 | Huaxia 19 | Henan Province 20220068 | 2022 |
| 620 | Jinyu 169 | Henan Province 20220069 | 2022 |
| 621 | Neimai 628 | Henan Province 20220070 | 2022 |
| 622 | Xinnong 25 | Henan Province 20220072 | 2022 |
| 623 | Yangguang 528 | Henan Province 20220073 | 2022 |
| 624 | Zhouyu 18 | Henan Province 20220075 | 2022 |
| 625 | Zhoufeng 173 | Henan Province 20220076 | 2022 |
| 626 | Yunong 606 | Henan Province 20220078 | 2022 |
| 627 | Yunong 810 | Henan Province 20220079 | 2022 |
| 628 | Shangnong 8 | Henan Province 20220080 | 2022 |
| 629 | Tunmai 296 | Henan Province 20220081 | 2022 |
| 630 | Nongyou 17 | Henan Province 20220082 | 2022 |
| 631 | Pingan 901 | Henan Province 20220084 | 2022 |
| 632 | Changmai 22 | Henan Province 20220086 | 2022 |
| 633 | Hemai 1707 | Henan Province 20220087 | 2022 |
| 634 | Jiyanmai 20 | Henan Province 20220088 | 2022 |
| 635 | Luomai 38 | Henan Province 20220089 | 2022 |
| 636 | Tianmin 355 | Henan Province 20220092 | 2022 |
| 637 | Xinmai 9389 | Henan Province 20220094 | 2022 |
| 638 | Zhengmai 186 | Henan Province 20220096 | 2022 |
| 639 | Zhengmai 188 | Henan Province 20220097 | 2022 |
| 640 | Zhengmai 0989 | Henan Province 20220099 | 2022 |
| 641 | Zhoumai 44 | Henan Province 20220100 | 2022 |
| 642 | Lianbang 3 | Henan Province 20220103 | 2022 |
| 643 | Nongmai 30 | Henan Province 20220104 | 2022 |
| 644 | Hemai 31 | Henan Province 20220107 | 2022 |
| 645 | Lantian 45 | Gansu Province 20220016 | 2022 |
| 646 | Hemai 317 | Shandong Province 20220014 | 2022 |
| 647 | Huangfanmai 58 | Shandong Province 20226021 | 2022 |
| 648 | Jimai 24 | Shandong Province 20226025 | 2022 |
| 649 | Nongke 1132 | Shaanxi Province 20220002 | 2022 |
| 650 | Xinbo 188 | Shaanxi Province 20220003 | 2022 |
| 651 | Xinong 136 | Shaanxi Province 20220004 | 2022 |
| 652 | Xichun 919 | Shaanxi Province 20220007 | 2022 |
| 653 | Xinong 186 | Shaanxi Province 20220008 | 2022 |
| 654 | Xinong 681 | Shaanxi Province 20220011 | 2022 |
| 655 | Xinong 865 | Shaanxi Province 20220013 | 2022 |
| 656 | Xinong 119 | Shaanxi Province 20220014 | 2022 |
| 657 | Xinong 256 | Shaanxi Province 20220015 | 2022 |
| 658 | Xinong 998 | Shaanxi Province 20220016 | 2022 |
| 659 | Yangzhi 171 | Shaanxi Province 20220018 | 2022 |
| 660 | Hanmai 9 | Shaanxi Province 20220021 | 2022 |
| 661 | Xinong 282 | Shaanxi Province 20220024 | 2022 |
| 662 | Weilong 181 | Shaanxi Province 20220025 | 2022 |
| 663 | Weilong 188 | Shaanxi Province 20220026 | 2022 |
| 664 | Xumai DH2 | Jiangsu Province 20220014 | 2022 |
| 665 | Fenghemai 693 | Jiangsu Province 20220017 | 2022 |
| 666 | Xumai 136 | Jiangsu Province 20220031 | 2022 |

| Number | Cultivar name | Provincial/National Certification | Year of release |
|---|---|---|---|
| 667 | Huamai 22 | Jiangsu Province 20220033 | 2022 |
| 668 | Bainong 5819 | National 2023031 | 2023 |
| 669 | Huaimai 58 | National 2023038 | 2023 |
| 670 | Kelin 201 | National 2023039 | 2023 |
| 671 | Kemai 2 | National 2023040 | 2023 |
| 672 | Linfanmai 3 | National 2023041 | 2023 |
| 673 | Luomai 68 | National 2023043 | 2023 |
| 674 | Ruihuamai 556 | National 2023046 | 2023 |
| 675 | Shangmai 189 | National 2023048 | 2023 |
| 676 | Shunmai 13 | National 2023049 | 2023 |
| 677 | Tianyikemai 8 | National 2023050 | 2023 |
| 678 | Xinong 1125 | National 2023053 | 2023 |
| 679 | Xinmai 58 | National 2023055 | 2023 |
| 680 | Xinzhi 6 | National 2023056 | 2023 |
| 681 | Xinliang 10 | National 2023057 | 2023 |
| 682 | Xumai 45 | National 2023058 | 2023 |
| 683 | Zhengmai 158 | National 2023060 | 2023 |
| 684 | Zhengmai 20 | National 2023062 | 2023 |
| 685 | Zhongdimai 668 | National 2023064 | 2023 |
| 686 | Zhongmai 698 | National 2023065 | 2023 |
| 687 | Zhongyu 1686 | National 2023066 | 2023 |
| 688 | Zhumai 586 | National 2023067 | 2023 |
| 689 | Anke 1804 | National 2023069 | 2023 |
| 690 | Fanmai 65 | National 2023071 | 2023 |
| 691 | Huamai 20 | National 2023072 | 2023 |
| 692 | Wansu 1232 | National 2023076 | 2023 |
| 693 | Guomai 303 | National 2023077 | 2023 |
| 694 | Xumai DH9 | National 2023080 | 2023 |
| 695 | Anke 1904 | National 2023085 | 2023 |
| 696 | Baomai 6 | National 2023087 | 2023 |
| 697 | Dingfu 196 | National 2023091 | 2023 |
| 698 | Guohemai 2 | National 2023092 | 2023 |
| 699 | Huawen 1 | National 2023095 | 2023 |
| 700 | Huaichuan 267 | National 2023096 | 2023 |
| 701 | Jinli 719 | National 2023099 | 2023 |
| 702 | Kelin 12 | National 2023101 | 2023 |
| 703 | Qimin 20 | National 2023104 | 2023 |
| 704 | Shunmai 20 | National 2023110 | 2023 |
| 705 | Taimai 701 | National 2023113 | 2023 |
| 706 | Tongzhou 9 | National 2023114 | 2023 |
| 707 | Xinong 112 | National 2023116 | 2023 |
| 708 | Xinong 115 | National 2023117 | 2023 |
| 709 | Bainong 607 | National 2023121 | 2023 |
| 710 | Xinkemai 170 | National 2023123 | 2023 |
| 711 | Xinzhi 716 | National 2023125 | 2023 |
| 712 | Xumai 2100 | National 2023127 | 2023 |
| 713 | Xumai 47 | National 2023128 | 2023 |
| 714 | Youfu 9 | National 2023129 | 2023 |

| Number | Cultivar name | Provincial/National Certification | Year of release |
|---|---|---|---|
| 715 | Zhimai 26 | National 2023132 | 2023 |
| 716 | Zhoumai 42 | National 2023134 | 2023 |
| 717 | Zhumai 126 | National 2023135 | 2023 |
| 718 | Zhuangmai 198 | National 2023136 | 2023 |
| 719 | Jinyongfeng 1 | National 2023145 | 2023 |
| 720 | Qimin 16 | National 2023146 | 2023 |
| 721 | Shannong 1695 | National 2023148 | 2023 |
| 722 | Qimin 24 | National 2023175 | 2023 |
| 723 | Mingmai 168 | Henan Province 2023001 | 2023 |
| 724 | Wenmai K3 | Henan Province 2023002 | 2023 |
| 725 | Wanmaiyihao | Henan Province 2023005 | 2023 |
| 726 | Zhouyuan 98 | Henan Province 2023006 | 2023 |
| 727 | Renmai 307 | Henan Province 2023009 | 2023 |
| 728 | Juke 8 | Henan Province 2023010 | 2023 |
| 729 | Fuyumai 201 | Henan Province 2023011 | 2023 |
| 730 | Kelin 11 | Henan Province 2023012 | 2023 |
| 731 | Yonglemai 836 | Henan Province 2023013 | 2023 |
| 732 | Tianhe 6 | Henan Province 2023014 | 2023 |
| 733 | Changmai 23 | Henan Province 2023017 | 2023 |
| 734 | Yangao 162 | Henan Province 2023022 | 2023 |
| 735 | Xinong 132 | Henan Province 2023023 | 2023 |
| 736 | Fanyumai 27 | Henan Province 2023024 | 2023 |
| 737 | Yanbo 505 | Henan Province 2023026 | 2023 |
| 738 | Yunong 269 | Henan Province 2023032 | 2023 |
| 739 | Jinmeng 3 | Henan Province 2023038 | 2023 |
| 740 | Wanmai 270 | Henan Province 2023040 | 2023 |
| 741 | Jiheimai 1 | Henan Province 2023042 | 2023 |
| 742 | Luoheimai 1 | Henan Province 2023043 | 2023 |
| 743 | Zhengmai 21 | Henan Province 2023048 | 2023 |
| 744 | Yunong 911 | Henan Province 2023049 | 2023 |
| 745 | Yunong 914 | Henan Province 2023050 | 2023 |
| 746 | Lunxuan 130 | Henan Province 2023054 | 2023 |
| 747 | Qiulemai 308 | Henan Province 2023057 | 2023 |
| 748 | Chengmai 829 | Henan Province 2023058 | 2023 |
| 749 | Jinmanliang 1 | Henan Province 2023059 | 2023 |
| 750 | Zhaofeng 26 | Henan Province 2023060 | 2023 |
| 751 | Zhaofeng 32 | Henan Province 2023061 | 2023 |
| 752 | Kaimai 1705 | Henan Province 2023062 | 2023 |
| 753 | Zhumai 156 | Henan Province 2023065 | 2023 |
| 754 | Luomai 198 | Henan Province 2023067 | 2023 |
| 755 | Huaichuan 109 | Henan Province 2023068 | 2023 |
| 756 | Huaichuan 938 | Henan Province 2023069 | 2023 |
| 757 | Xuke 772 | Henan Province 2023070 | 2023 |
| 758 | Jinma 1 | Henan Province 2023071 | 2023 |
| 759 | Mengmai 188 | Henan Province 2023072 | 2023 |
| 760 | Neinongke 203 | Henan Province 2023073 | 2023 |
| 761 | Yuhong 99 | Henan Province 2023074 | 2023 |
| 762 | Yuyuan 8 | Henan Province 2023076 | 2023 |

| Number | Cultivar name | Provincial/National Certification | Year of release |
|---|---|---|---|
| 763 | Zhongjie 3 | Henan Province 2023078 | 2023 |
| 764 | Zhouqun 6 | Henan Province 2023079 | 2023 |
| 765 | Zhouqun 58 | Henan Province 2023080 | 2023 |
| 766 | Zhouqun 9 | Henan Province 2023081 | 2023 |
| 767 | Zhouyu 801 | Henan Province 2023082 | 2023 |
| 768 | Huaguan 181 | Henan Province 2023083 | 2023 |
| 769 | Huaguan 184 | Henan Province 2023084 | 2023 |
| 770 | Huaguan 174 | Henan Province 2023087 | 2023 |
| 771 | Yunong 521 | Henan Province 2023088 | 2023 |
| 772 | Jinli 826 | Henan Province 2023091 | 2023 |
| 773 | Tianning 58 | Henan Province 2023092 | 2023 |
| 774 | Wenmai 802 | Henan Province 2023093 | 2023 |
| 775 | Luomai 77 | Henan Province 2023095 | 2023 |
| 776 | Wan 1393 | Henan Province 2023097 | 2023 |
| 777 | Xinmai 57 | Henan Province 2023098 | 2023 |
| 778 | Anmai 19 | Henan Province 2023100 | 2023 |
| 779 | Bainong 8399 | Henan Province 2023101 | 2023 |
| 780 | Changmai 169 | Henan Province 2023102 | 2023 |
| 781 | Luomai 44 | Henan Province 2023103 | 2023 |
| 782 | Shengmaiyuan 789 | Henan Province 2023106 | 2023 |
| 783 | Shengmaiyuan 3268 | Henan Province 2023107 | 2023 |
| 784 | Xinyu 183 | Henan Province 2023108 | 2023 |
| 785 | Tongzhou 5 | Henan Province 2023109 | 2023 |
| 786 | Tongzhou 6 | Henan Province 2023110 | 2023 |
| 787 | Jinkemai 11 | Henan Province 2023111 | 2023 |
| 788 | Hemai 66 | Henan Province 2023112 | 2023 |
| 789 | Shenzhoumai 227 | Henan Province 2023113 | 2023 |
| 790 | Delichangmai 2 | Henan Province 2023114 | 2023 |
| 791 | Hemai 58 | Henan Province 2023115 | 2023 |
| 792 | Qihang 66 | Henan Province 2023116 | 2023 |
| 793 | Zhengke 26 | Henan Province 2023117 | 2023 |
| 794 | Delichangmai 4 | Henan Province 2023119 | 2023 |
| 795 | Shenzhoumai 222 | Henan Province 2023120 | 2023 |
| 796 | Hongyumai 2 | Henan Province 2023121 | 2023 |
| 797 | Jiashuo 99 | Henan Province 2023122 | 2023 |
| 798 | Hainamai 211 | Henan Province 2023123 | 2023 |
| 799 | Chenbo 985 | Henan Province 2023124 | 2023 |
| 800 | Shuangdemai 9 | Henan Province 2023126 | 2023 |
| 801 | Niumai 168 | Henan Province 2023127 | 2023 |
| 802 | Yishuomai 139 | Henan Province 2023128 | 2023 |
| 803 | Caimai 208 | Henan Province 2023129 | 2023 |
| 804 | Huamai 128 | Hubei Province 20230007 | 2023 |
| 805 | Emai 016 | Hubei Province 20230010 | 2023 |
| 806 | Huamai 1820 | Hubei Province 20230014 | 2023 |
| 807 | Yingpoaifeng 8 | Hebei Province 20238003 | 2023 |
| 808 | Jiyuan 3949 | Hebei Province 20238008 | 2023 |
| 809 | Xinong 962 | Shaanxi Province 20230001 | 2023 |
| 810 | Xianmai 341 | Shaanxi Province 20230004 | 2023 |

| Number | Cultivar name | Provincial/National Certification | Year of release |
|---|---|---|---|
| 811 | Xinong 226 | Shaanxi Province 20230005 | 2023 |
| 812 | Xinong 1165 | Shaanxi Province 20230010 | 2023 |
| 813 | Weilong 323 | Shaanxi Province 20230011 | 2023 |
| 814 | Baoyan 12 | Shaanxi Province 20230012 | 2023 |
| 815 | Qiushi 369 | Shaanxi Province 20230014 | 2023 |
| 816 | Xinong 1699 | Shaanxi Province 20230015 | 2023 |
| 817 | Xinong 5812 | Shaanxi Province 20230016 | 2023 |
| 818 | Xinong 591 | Shaanxi Province 20230017 | 2023 |
| 819 | Weilong 396 | Shaanxi Province 20230025 | 2023 |
| 820 | Lvyu 1102 | Anhui Province 2023L002 | 2023 |
| 821 | Xinshiji 156 | Anhui Province 2023L007 | 2023 |
| 822 | Yutianmai 1891 | Anhui Province 2023L010 | 2023 |
| 823 | Wannong 211 | Anhui Province 2023L018 | 2023 |
| 824 | Fumai 20 | Anhui Province 2023L019 | 2023 |
| 825 | Fumai 14 | Anhui Province 2023L020 | 2023 |
| 826 | Youxinmai 288 | Anhui Province 2023L023 | 2023 |
| 827 | Mengmai 313 | Anhui Province 2023L024 | 2023 |
| 828 | Fumai 19 | Anhui Province 2023T001 | 2023 |
| 829 | Su 6165 | Anhui Province 2023T005 | 2023 |
| 830 | Wansu 1235 | Anhui Province 2023T006 | 2023 |
| 831 | Quanmai 838 | Anhui Province 2023T009 | 2023 |
| 832 | Lvyu 18 | Anhui Province 2023T011 | 2023 |
| 833 | Dadi 202 | Anhui Province 2023T014 | 2023 |
| 834 | Sui 1808 | Anhui Province 2023T016 | 2023 |
| 835 | Wankenmai 209 | Anhui Province 2023T018 | 2023 |
| 836 | Guomai 19 | Anhui Province 2023T020 | 2023 |
| 837 | Qinglin 139 | Anhui Province 2023T021 | 2023 |
| 838 | Annong 197 | Anhui Province 2023T024 | 2023 |
| 839 | Jinke 788 | Anhui Province 2023T028 | 2023 |

**Table S2. Summary of the HiFi reads used in developing Zhou8425B genome assembly**

| Reads number | Base number (Gb) | Mean length (kb) | N50 (kb) | Depth (X) |
|---|---|---|---|---|
| 43,733,624 | 728.84 | 16.66 | 16.58 | 48 |

**Table S3. Summary of the Hi-C reads used in developing Zhou8425B genome assembly**

| Clean reads (pair) | Clean bases (Gb) | Depth (X) | Alignable paired | Uniquely paired |
|---|---|---|---|---|
| 6,004,887,544 | 1,801.46 | 124 | 4,289,278,542 (71.42%) | 1,405,265,150 (23.40%) |

**Table S4. Statistics of the assembled Zhou8425B chromosomes**

| Chromosome | Contig number | Chromosome length (Mb) | Size of assembled subgenome (Gb) |
|---|---|---|---|
| Chr1A | 15 | 604.64 | |
| Chr2A | 25 | 789.23 | |
| Chr3A | 11 | 759.88 | |
| Chr4A | 42 | 767.79 | 5.01 (Subgenome A) |
| Chr5A | 12 | 717.45 | |
| Chr6A | 10 | 625.04 | |
| Chr7A | 17 | 754.24 | |
| Chr1RS-1BL | 40 | 785.82 | |
| Chr2B | 46 | 820.21 | |
| Chr3B | 85 | 866.06 | |
| Chr4B | 55 | 708.42 | 5.43 (Subgenome B) |
| Chr5B | 75 | 735.10 | |
| Chr6B | 81 | 745.47 | |
| Chr7B | 71 | 765.69 | |
| Chr1D | 5 | 500.93 | |
| Chr2D | 10 | 663.77 | |
| Chr3D | 14 | 639.59 | |
| Chr4D | 6 | 532.02 | 4.08 (Subgenome D) |
| Chr5D | 15 | 585.45 | |
| Chr6D | 6 | 504.40 | |
| Chr7D | 12 | 652.21 | |
| Total | 653 | 14,523.41 | 14.52 |

**Table S5. The assembled centromeres of Zhou8425B chromosomes**

| Chromosome | Centromere start (Mb) | Centromere end (Mb) | Contig number* |
|---|---|---|---|
| Chr1A | 213.10 | 222.80 | 1 |
| Chr1B | 314.20 | 320.20 | 1 |
| Chr1D | 175.50 | 177.40 | 1 |
| Chr2A | 341.30 | 349.50 | 1 |
| Chr2B | 365.20 | 370.80 | 1 |
| Chr2D | 273.70 | 278.10 | 1 |
| Chr3A | 325.00 | 332.40 | 1 |
| Chr3B | 368.20 | 375.00 | 1 |
| Chr3D | 261.20 | 267.40 | 1 |
| Chr4A | 289.30 | 298.70 | 1 |
| Chr4B | 284.20 | 290.50 | 1 |
| Chr4D | 213.40 | 216.40 | 1 |
| Chr5A | 246.90 | 255.40 | 1 |
| Chr5B | 228.60 | 236.20 | 1 |
| Chr5D | 195.00 | 201.00 | 1 |
| Chr6A | 289.20 | 295.10 | 1 |
| Chr6B | 343.80 | 351.60 | 1 |
| Chr6D | 242.90 | 244.70 | 1 |
| Chr7A | 369.60 | 373.90 | 1 |
| Chr7B | 312.70 | 317.40 | 1 |
| Chr7D | 345.20 | 351.20 | 1 |

*The number of contigs used to assemble the centromere.

**Table S6. Summary of the transcriptome sequencing data obtained in this study**

| Plant sample | Q20 (%) | Q30 (%) | GC content (%) | Clean reads | Clean base (Gb) | Mapping rate (%)* |
|---|---|---|---|---|---|---|
| 10-day seedling leaf (SL) | 96.82 | 91.82 | 55.97 | 91,699,548 | 13.54 | 93.67 |
| 10-day seedling root (SR) | 97.04 | 92.26 | 54.05 | 81,386,040 | 11.99 | 92.85 |
| Heading-stage leaf (HL) | 97.34 | 93.08 | 57.52 | 99,556,542 | 14.65 | 93.31 |
| Heading-stage root (HR) | 97.18 | 92.66 | 55.25 | 94,370,092 | 13.92 | 92.75 |
| Heading-stage stem (HS) | 96.84 | 90.77 | 55.71 | 80,428,908 | 11.91 | 93.82 |
| Heading-stage stem node (HSN) | 97.07 | 92.38 | 54.86 | 91,733,766 | 13.56 | 93.23 |
| Heading-stage ineffective tiller (HIT) | 97.36 | 93.10 | 55.32 | 104,121,440 | 15.34 | 94.33 |
| Heading-stage ineffective tiller bud (HITB) | 97.07 | 92.34 | 55.76 | 89,202,258 | 13.16 | 93.05 |
| Immature spike (1 cm) (IS1) | 97.06 | 92.37 | 53.87 | 95,026,038 | 14.05 | 92.35 |
| Immature spike (2-3 cm) (IS2) | 97.02 | 92.26 | 54.53 | 85,171,830 | 12.55 | 92.11 |
| 5 day developing grain (5DG) | 97.08 | 92.45 | 56.11 | 92,733,680 | 13.71 | 93.75 |
| 15 day developing grain (15DG) | 96.55 | 91.61 | 54.69 | 88,494,498 | 12.97 | 91.72 |

*Calculated by mapping to the genome assembly of Zhou8425B.

**Table S7. Functional annotation of the 106,940 HC protein-coding genes of Zhou8425B**

| Database | Number | Percentage (%) |
|---|---|---|
| Annotated by NR | 105,638 | 98.78 |
| Annotated by Swiss Prot | 71,352 | 66.72 |
| Annotated by KEGG | 38,747 | 36.23 |
| Annotated by GO | 46,238 | 43.24 |
| Annotated by eggNOG | 94,686 | 88.54 |
| Total annotated genes | 105,649 | 98.79 |
| Not Annotated | 1,291 | 1.21 |

**Table S8. Summary of non-coding RNA genes annotated for Zhou8425B genome assembly**

| Class | Number | Average length (bp) | Total length (Mb) |
|---|---|---|---|
| rRNA | 26,237 | 1,219 | 31.99 |
| tRNA | 51,883 | 75 | 3.92 |
| miRNA | 39,521 | 128 | 5.06 |
| snRNA | 1,147 | 143,109 | 0.16 |
| snoRNA | 3,973 | 1,219 | 0.43 |

**Table S9. Statistics of repeat elements annotated for Zhou8425B genome assembly**

| Class | Super family | Family | Count | Size (bp) | Percentage* |
|---|---|---|---|---|---|
| Class I: | LTR element | Gypsy | 2,004,608 | 6,443,818,348 | 43.66 |
| | | Copia | 789,814 | 2,362,913,823 | 16.01 |
| | | Unclassified | 403,413 | 709,996,845 | 4.81 |
| | Non-LTR element | LINE | 104,207 | 78,814,310 | 0.53 |
| | | SINE | 6,513 | 1,245,977 | 0.01 |
| Class II: | DNA TE | CACTA | 1,690,749 | 2,219,542,183 | 15.04 |
| | | Mutator | 290,234 | 128,894,277 | 0.87 |
| | | Unclassified TIRs | 100,809 | 26,337,731 | 0.18 |
| | | PIF/Harbinger | 175,472 | 77,920,827 | 0.53 |
| | | Tcl/Mariner | 485,710 | 158,518,870 | 1.07 |
| | | Unclassified | 12,968 | 9,287,897 | 0.06 |
| | | hAT | 105,924 | 41,012,410 | 0.28 |
| | Helitron | | 248,260 | 108,107,229 | 0.73 |
| Others | | | 925,945 | 544,366,145 | 3.69 |
| Total | | | 7,344,626 | 12,910,776,872 | 87.53 |

*Calculated based on the 14.75 Gb genome assembly of Zhou8425B.

**Table S10. Mapping result of HiFi reads to Zhou8425B genome assembly**

| Reads number | Reads properly mapped | Mapping ratio |
|---|---|---|
| 43,318,145 | 43,302,959 | 99.96% |

**Table S11. Mapping result of Illumina reads to Zhou8425B assembled chromosomes**

| Chromosome | QV | Nucleotide accuracy (%) |
|---|---|---|
| Chr1A | 48.4624 | 99.998575 |
| Chr1B | 46.8787 | 99.997948 |
| Chr1D | 47.0801 | 99.998041 |
| Chr2A | 47.9779 | 99.998407 |
| Chr2B | 47.355 | 99.998161 |
| Chr2D | 45.0251 | 99.996856 |
| Chr3A | 48.2913 | 99.998518 |
| Chr3B | 46.9326 | 99.997974 |
| Chr3D | 47.1127 | 99.998056 |
| Chr4A | 47.9372 | 99.998392 |
| Chr4B | 47.247 | 99.998115 |
| Chr4D | 47.1387 | 99.998067 |
| Chr5A | 48.3094 | 99.998524 |
| Chr5B | 47.0763 | 99.998039 |
| Chr5D | 47.1477 | 99.998071 |
| Chr6A | 48.2653 | 99.998509 |
| Chr6B | 47.1973 | 99.998093 |
| Chr6D | 47.0025 | 99.998006 |
| Chr7A | 48.0185 | 99.998422 |
| Chr7B | 46.9918 | 99.998001 |
| Chr7D | 46.86 | 99.997939 |

**Table S12. BUSCO evaluation result of Zhou8425B genome assembly**

| Complete and single-copy | Complete duplicated | Fragmented | BUSCO completeness | Total BUSCO genes searched |
|---|---|---|---|---|
| 133 (2.7%) | 4733 (96.7%) | 3 (0.06%) | 4866 (99.4%) | 4,896 (BUSCO version: 5.1.3; Lineage dataset: poales_odb10) |

**Table S13. Summary of gluten gene loci and contigs identified in Zhou8425B genome assembly** (shown as separate excel file)

**Table S14. List of 922 genes co-expressed with TraesZ8425B1RS01G036200** (shown as separate excel file)

**Table S15. SNP markers significantly associated with flag leaf disease severity (FLDS) detected in two years in Xindu (XD) city of Sichuan province by GWAS analysis** (shown as separate excel file)

**Table S16. SNP markers significantly associated with flag leaf disease severity (FLDS) detected in three years in Pixian (PX) city of Sichuan province by GWAS analysis** (shown as separate excel file)

**Table S17. Chi-squared test of segregation displayed by the 198 $F_{2:3}$ lines of Xinhuamai 818/Yumai 1 population in response to yellow rust infections**

| Plant material | Number of plants | Flag leaf disease severity (FLDS) | | |
|---|---|---|---|---|
| | | Homozygous and resistant (FLDS < 45%) | Heterozygous and resistant (20% < FLDS < 85%) | Homozygous and susceptible (FLDS > 55%) |
| Xinhuamai 818 | 30 | 30 | | |
| Yumai 1 | 30 | | | 30 |
| $F_{2:3}$* | 199 | 42 | 101 | 56 |

*The segregation ratio of $F_{2:3}$ lines conformed with a 1:2:1 ratio ($\chi^2 = 2.02 < \chi^2_{0.05, 2} = 5.99$).

**Table S18. List of ten polymorphic DNA markers developed for fine mapping of *YrZH3BS***

| Name | Primer (5′- 3′) | Fragment size (bp) amplified from resistant/susceptible lines | Physical location on 3BS (Mb) |
|---|---|---|---|
| M174 | F: CAGCACCTGCCCGACAATAA<br>R: CTCCGTGATAGGGAAAGCCC | 193/181 | 7.19 |
| M199 | F: CCAAACCCAAAGCCAAGTCG<br>R: GCGAGTCACCGAATTGAACG | 124/118 | 8.53 |
| M207 | F: TTCACCGCTCAGGGTAGTAAG<br>R: AGGTACACCCCGTTCTCGAT | 226/234 | 8.79 |
| M287 | F: GGGTGAGGATGAGGAACCCT<br>R: AGAGCTCTGCGAAGAAACCA | 108/78 | 12.32 |
| M328 | F: CCGGCAATGTCAAGGACAAG<br>R: ATCGAAACGTGGAAAGCAACAG | 273/228 | 13.25 |
| M311 | F: TGTGCTAAGAGCCTGAGGGA<br>R: CCAACGCACTCTACCACAGT | 177/152 | 14.16 |
| M344 | F: CACCAGTAGGCGCTCTCATC<br>R: CCCTGTATTCTCAGCACCCC | 146/177 | 16.44 |
| M392 | F: TGATAATCCTGTTGCTCTTTCCGA<br>R: TGTGCAGTATGTTCGATTAGGTTC | 90/252 | 18.86 |
| M409 | F: TGATCCATGTATGTGCTGCTTCT<br>R: GCGAGATATATACAGCGGCCAG | 95/120 | 20.35 |

**Table S19. List of 212 common wheat cultivars and information on their flag leaf disease severity, yield related trait (SNPP, GNPS, TGW, and GY), and adult plant resistance gene data** (shown as separate excel file)

**Table S20. List of PCR primers used for detecting 17 agronomic trait genes in Zhou8425B and the common wheat lines used during breeding of Zhou8425B**

| Trait gene | Primers (5′- 3′) | Reference |
|---|---|---|
| *Rht1* | F: TCTCCTCCCTCCCCACCCCAAC<br>R: CATCCCCATGGCCATCTCGAGCTA | Ellis M, Spielmeyer W, Gale K, Rebetzke G, Richards R. "Perfect" markers for the *Rht-B1b* and *Rht-D1b* dwarfing genes in wheat. Theor Appl Genet. 2002; 105: 1038-1042. |
| *Rht2* | F: CGCGCAATTATTGGCCAGAGATAG<br>R: CCCCATGGCCATCTCGAGCTGCTA | Ellis M, Spielmeyer W, Gale K, Rebetzke G, Richards R (2002) "Perfect" markers for the *Rht-B1b* and *Rht-D1b* dwarfing genes in wheat. Theor Appl Genet. 105: 1038-1042. |
| *Rht8* | F: CTTGACGAGCTTGGAAATGG<br>R: GCAACAAGTGCTTCTGTCGT | Asplund L, Leino MW, Hagenblad J. Allelic variation at the *Rht8* locus in a 19th century wheat collection. Scientific World J. 2012; 2012: 385610. |
| *Rht24* | F: TCGCAACTTTTGCTAGGCAAT<br>R: CAGGAGCAATCTACAGCAGAAAG | Tian X, Xia X, Xu D, Liu Y, Xie L, Hassan MA, Song J, Li F, Wang D, Zhang Y, Hao Y, Li G, Chu C, He Z, Cao S. *Rht24b*, an ancient variation of *TaGA2ox-A9*, reduces plant height without yield penalty in wheat. New Phytol. 2022; 233: 738-750. |
| *Ppd-D1a* | F: ACGCCTCCCACTACACTG<br>R: GTTGGTTCAAACAGAGAGC | Beales J, Turner A, Griffiths S. A pseudoresponse regulator is misexpressed in the photoperiod insensitive *Ppdl-D1a* mutant of wheat (*Triticum aestivum* L.). Theor Appl Genet. 2007; 115: 721-733. |
| *YrZH84* | F: TCAAATGATTTCAGGTAACCACTA<br>R: TTCCTGATCCCACCAAACAT | Li ZF, Zheng TC, He ZH, Li GQ, Xu SC, Li XP, Yang GY, Singh RP, Xia XC. Molecular tagging of stripe rust resistance gene *YrZH84* in Chinese wheat line Zhou 8425B. Theor Appl Genet. 2006; 112: 1198-1103. |
| *YrZH22* | F: CGGCCAAATATGAGACTGCC<br>R: AATGCGGTGAATGGAAGACG | Wang Y, Xie J, Zhang H, Guo B, Ning S, Chen Y, Lu P, Wu Q, Li M, Zhang D, Guo G, Zhang Y, Liu D, Zou S, Tang J, Zhao H, Wang X, Li J, Yang W, Cao T, Yin G, Liu Z. Mapping stripe rust resistance gene *YrZH22* in Chinese wheat cultivar Zhoumai 22 by bulked segregant RNA-Seq (BSR-Seq) and comparative genomics analyses. Theor Appl Genet. 2017; 130: 2191-2201. |
| *Yr3BS* | F: CCGGCAATGTCAAGGACAAG<br>R: ATCGAAACGTGGAAAGCAACAG | Developed in this study. |
| *LrZH84* | F: GTCTGCAAACTTGAAGGAAG<br>R: GCAGATTTCAGTTCATCCTC | Zhou Y, Xia X, He Z, Li X, Li Z, Liu D. Fine mapping of leaf rust resistance gene *LrZH84* using expressed sequence tag and sequence-tagged site markers, and allelism with other genes on wheat chromosome 1B. Phytopathology. 2013; 103: 169-74. |
| *Lr13* | F: TGCACATGGATATGCGGAGA<br>R: GCTTTGAGTTGCCCATGCTC | Hewitt T, Zhang J, Huang L, Upadhyaya N, Li J, Park R, Hoxha S, McIntosh R, Lagudah E, Zhang P. Wheat leaf rust resistance gene *Lr13* is a specific *Ne2* allele for hybrid necrosis. Mol Plant. 2021; 14: 1025-1028. |
| *TaCwi-A1* | FAM: GATGAAACAAGTTAGTCCGGTAC<br>HEX: GATGAAACAAGTTAGTCCGGTAA<br>R: CTCCAACTGAACTGGTGCAA | Ma D, Yan J, He Z, Wu L, Xia X. Characterization of a cell wall invertase gene *TaCwi-A1* on common wheat chromosome 2A and development |

| | | |
|---|---|---|
| | | of functional markers. Mol Breeding. 2012; 29: 43-52. |
| *TaSus1-7A* | FAM: GATTTGATCCATGCCCTCTC<br>HEX: GATTTGATCCATGCCCTCTT<br>R: CTGTCGTTCAACATCATTGTCTG | Hou J, Jiang Q, Hao C, Wang Y, Zhang H, Zhang X. Global selection on sucrose synthase haplotypes during a century of wheat breeding. Plant Physiol. 2014; 164: 1918-1929. |
| *TaSus1-7B* | FAM: CAATTGCTTATGTTCTGTTGTATGG<br>HEX: CAATTGCTTATGTTCTGTTGTACAT<br>R: ATGGTTATGCTTGAATGGAAGAGC | Hou J, Jiang Q, Hao C, Wang Y, Zhang H, Zhang X. Global selection on sucrose synthase haplotypes during a century of wheat breeding. Plant Physiol. 2014; 164: 1918-1929. |
| *TaSus2-2A* | FAM: AGCAATGGGGGAGACTGCTGGAGG<br>HEX: AGCAATGGGGGAGACTGCTGGAGA<br>R: GCTGTGGATGCGGCTCAGGGCGCG | Hou J, Jiang Q, Hao C, Wang Y, Zhang H, Zhang X. Global selection on sucrose synthase haplotypes during a century of wheat breeding. Plant Physiol. 2014; 164: 1918-1929. |
| *TaGS-D1* | FAM: GCCAAGAAATGTCGCTCTCAG<br>HEX: GCCAAGAAATGTCGCTCTCAT<br>R: CAAGAATTTTGGGACGGAGGGA | Rasheed A, Wen W, Gao F, Zhai S, Jin H, Liu J, Guo Q, Zhang Y, Dreisigacker S, Xia X, He Z. Development and validation of KASP assays for genes underpinning key economic traits in bread wheat. Theor Appl Genet. 2016; 129: 1843-1860. |
| *TaGS5-A1* | FAM: GTGCAATCTTGGACAAACATCAG<br>HEX: GTGCAATCTTGGACAAACATCAT<br>R: AGTGCTTTGTCAACAACAGATGC | Ma L, Li T, Hao C, Wang Y, Chen X, and Zhang X. *TaGS5-3A*, a grain size gene selected during wheat improvement for larger kernel and yield. Plant Biotechnol J. 2016; 14: 1269-1280. |
| *TaGW2-6B* | FAM: GTTGGTGTCATTTGTAAAGCCCA<br>HEX: GTTGGTGTCATTTGTAAAGCCCC<br>R: AGGCTTGTCAAAACGTGGGGTCC | Qin L, Hao C, Hou J, Wang Y, Li T, Wang L, Ma Z, Zhang X. Homologous haplotypes, expression, genetic effects and geographic distribution of the wheat yield gene *TaGW2*. BMC Plant Biol. 2014; 14: 107. |
